# Kernel-based LFP estimation in detailed large-scale spiking network model of mouse visual cortex

**DOI:** 10.1101/2024.11.29.626029

**Authors:** Nicolò Meneghetti, Atle E. Rimehaug, Gaute T. Einevoll, Alberto Mazzoni, Torbjørn V. Ness

## Abstract

Simulations of large-scale neural activity are powerful tools for investigating neural networks. Calculating measurable brain signals like local field potentials (LFPs) bridges the gap between model predictions and experimental observations. However, accurately simulating LFPs from large-scale models has traditionally required highly detailed multicompartmental neuron models, posing significant computational challenges. Here, we demonstrate that a kernel-based method can efficiently and accurately estimate LFPs in a state-of-the-art multicompartmental model of the mouse primary visual cortex (V1). Beyond its computational efficiency, the kernel method aids analysis by disentangling contributions of individual neuronal populations to the LFP. Using this approach, we found that LFPs in the V1 model were dominated by external synaptic inputs, with local synaptic activity playing a minimal role. Our findings establish the kernel method as a powerful tool for LFP estimation in large-scale network models and for uncovering the synaptic mechanisms underlying brain signals.

## Introduction

Extracellular potentials are measurable electric signals generated by the ion flow during neuronal activity ^1–3^. The low-frequency component of extracellular potentials, typically below a few hundred Hz, is referred to as the local field potential (LFP). The LFP primarily reflects the integrated synaptic activity of large populations of neurons, covering spatial scales from several hundred micrometers to a few millimeters ^1,4–8^.

Because of the ability to capture neural dynamics at a mesoscopic scale, LFPs have become an essential tool for studying neuronal function. Advances in multicontact electrodes for high-density laminar recordings ^9–12^ have further heightened interest in LFPs, providing improved spatial resolution and coverage. Additionally, the temporal stability of LFP recordings makes them attractive for applications in brain-computer interfaces and neuroprosthetics ^13,14^, where sustained and reliable signals are crucial.

The versatility of LFPs is evident in their widespread use across various domains of neuroscience. LFPs have been instrumental in uncovering network mechanisms underlying sensory processing ^15–23^, motor planning ^24–29^, and higher cognitive processes including attention, memory, and perception ^30–35^. Additionally, LFPs have emerged as important biomarkers for various neurological disorders ^36^, including epilepsy ^37,38^, glioma ^39–41^, schizophrenia ^42,43^, migraine ^44^, Alzheimer’s disease, ^45^ and Parkinson’s disease ^46–48^.

The growing focus on local field potentials as a window into neural activity at the systems level has spurred the development of computational models to better understand their biophysical origins ^2,6,49–61^. These efforts typically rely on multicompartmental (MC) neuron models to simulate transmembrane currents, coupled with volume conductor (VC) theory to predict extracellular potentials. While this approach is biophysically well-founded, it can be computationally expensive, particularly for large-scale network models.

As a result, researchers have developed more efficient methods for estimating LFPs. One notable approach is the hybrid scheme^53^, which separates network spiking activity simulations from the subsequent LFP predictions. Spikes, simulated in point-neuron networks, are stored and later used to drive synaptic currents in unconnected MC neuron models, which in turn predict transmembrane currents and ultimately LFPs^62^.

A simpler method^63^ approximated LFPs as weighted, time-shifted sums of excitatory and inhibitory synaptic currents extracted from computationally efficient point-neuron simulations.

Other studies have exploited the near-linear relationship between action potential timing and extracellular potentials to estimate LFPs. This can be achieved by convolving presynaptic firing rates with linear filters, or ‘kernels’, which capture the spatiotemporal dynamics of synaptic inputs. Each of these kernels represents the averaged causal spike-signal impulse response function for pairs of pre- and post-synaptic populations, thus capturing the average postsynaptic LFP contribution given an action potential in the presynaptic population. The overall LFP can be estimated by convolving the firing rate of each presynaptic population with its corresponding population kernel, and summing all of these LFP contributions.

(Hagen et al., 2016)^53^ applied this kernel-based approach to a multilayer point-neuron network, demonstrating that LFPs can be predicted by convolving precomputed spatiotemporal kernels with presynaptic firing rates. The kernels were defined as the extracellular response to the synchronous activation of all neurons in a presynaptic population, normalized by the size of the presynaptic population. Although the kernel method was demonstrated to be accurate, and efficient once the kernels were known, estimating the kernels was highly computationally demanding.

In contrast, Teleńczuk et al. (2020)^64^ took a different approach, focusing on the experimental estimation of LFP kernels. Instead of precomputing them from simulations, they fitted monosynaptic extracellular responses to predefined spatiotemporal shape functions for excitatory and inhibitory inputs. These experimentally derived spike-trigger averaged LFP kernels were then used to approximate LFP signals, by convolving them with firing rates from point-neuron network simulations. However, kernels measured from spike-triggered averages are potentially troubled by correlations and network dynamics ^53^, and it is not a priori clear if such measured LFP kernels are transferable to other brain areas or species from where they were measured.

More recently, (Hagen et al., 2022)^55^ introduced a novel method that directly derived LFP kernels from the biophysical properties of neuronal networks, based on the spatial distribution of cells and synapses, conduction delays, synaptic dynamics, and firing rates.

In essence, this framework leveraged a single, biophysically detailed cell simulation to predict the population kernels. This was achieved by first simulating the transmembrane currents of a single postsynaptic neuron in response to conductance-based synaptic inputs, allowing this to represent the population-averaged membrane currents following presynaptic activation. Factors such as the spatial extent of the population and variability in synaptic parameters were then incorporated through a series of linear convolutions applied across both spatial and temporal domains before the LFP kernel was calculated. This method significantly enhanced the kernel approach’s efficiency and versatility, enabling accurate and efficient calculation of LFP kernels on standard computational platforms, such as common laptops ^65^.

The kernel method has shown a lot of promise for enabling efficient and accurate LFP estimation from neural simulations. The method has, however, only been validated for simple networks ^65^, and remains to be tested in more biologically detailed networks that feature multiple neuronal populations, intricate connectivity patterns, variable synaptic strengths, and fluctuating external inputs.

Here, we investigated this aspect by testing the kernel method proposed by (Hagen et al., 2022) ^55^ against a computational model of the mouse primary visual cortex (V1) developed by the Allen Institute, which features an unprecedented level of biological detail ^66^. Briefly, the V1 network comprises >50,000 multicompartmental Hodgkin–Huxley neurons belonging to 17 cell types (4 excitatory, 4 parvalbumin-, 4 somatostatin-, and 5 5- hydroxytryptamine- expressing interneurons) distributed in six cortical layers. Recurrent connection probabilities depend on intersomatic distance as well as neurons’ functional preferences, such as their direction tuning, with synaptic strength following an orientation-dependent like-to-like rule. Additionally, the model receives sensory inputs from three different external sources: (i) experimentally recorded afferent activity from the lateral geniculate nucleus (LGN); (ii) experimentally recorded feedback from the lateromedial visual cortex (we employed the version presented by (Rimehaug et al., 2023)^59^, distinguished by its novel integration of experimentally recorded feedback from the higher lateromedial (LM) visual area); and (iii) Poisson background spiking activity representing stimulus-independent continuous influence of the rest of the brain. To simplify analysis and effectively evaluate the kernel method, we first constructed a simplified version of the V1 network, retaining the full biological complexity of the original model but focusing only on neurons with somata located in layer 2/3 (L2/3). This approach allowed us to isolate a subset of the network where the kernel method could be rigorously tested.

The simulated firing rates and membrane potentials in the L2/3 model exhibited high levels of non-stationarity, which we found to significantly influence LFP generation. By incorporating these fluctuations into the kernel estimation, we demonstrated that the kernel method provides accurate LFP predictions for this highly detailed network model.

We compared the effectiveness of the kernel method to that of a proxy based on a linear combination of synaptic currents extracted from point neuron model simulations, as in (Mazzoni et al. 2015). While this alternative approach worked, it proved impractical in most cases due to the large number of free parameters involved.

We also extended the application of the kernel method to the full-column V1 model, which includes all cortical layers. While the method produced reliable LFP estimates in L2/3 and L4, its performance in L5 was notably less consistent. This discrepancy, however, allowed us to identify a likely artifact in the V1 model itself, which appeared to contribute to the observed inaccuracies. Thus, the kernel method also served as a diagnostic tool, revealing a putative model artifact that could otherwise have remained undetected.

Beyond its computational efficiency for LFP estimation, the kernel method also provides a valuable means of disentangling the contributions of individual neuronal populations to the overall LFP. Leveraging this opportunity, we found that the LFP in this model was primarily driven by synaptic input onto pyramidal cells from external sources, namely the lateromedial visual area in L2/3 and the lateral geniculate nucleus in L4. In contrast, local synaptic inputs from excitatory or inhibitory populations played only a negligible role in shaping the overall LFP.

We here used the large-scale highly biophysically detailed Allen Institute’s V1 model, which requires high-performance computing facilities, potentially limiting its accessibility within the neuroscience community. We demonstrate that accurate LFP estimation can be done through the kernel-based framework at a fraction of the computational cost, thereby facilitating detailed, accessible, and versatile LFP estimation from a wide range of network models at different levels of abstraction ^57^.

This study is structured as follows. We begin by detailing the network properties of the layer 2/3 V1 model and the kernel-based method for estimating LFPs in these upper cortical layers. We then present our novel kernel method, which accounts for network non-stationarity by incorporating fluctuations in presynaptic firing rates and postsynaptic membrane potentials during kernel estimation. Next, we systematically disentangle the contributions of external and internal synaptic inputs to the LFPs generated by cells in layers 2/3. Further, we describe two distinct approaches for estimating LFPs from a point-neuron network: (i) convolution with precomputed kernels, and (ii) identifying the optimal linear combination of synaptic currents that best captures the LFP time course, following a method similar to that of ^63^. Lastly, we extend the applicability of the kernel-based approach to layer 4 and discuss some of its limitations when applied to layer 5.

## Results

### LFPs from detailed large-scale L2/3 network model

We first investigated the LFPs generated by neurons with somata positioned in a single cortical layer, layer 2/3 (L2/3, see Methods), across the whole cortical depth, upon presentation of simple visual stimuli. To this end, we built on the mouse V1 multi-compartmental full-column model developed by the Allen Institute ^66^, specifically the version presented by (Rimehaug et al., 2023)^59^, mainly distinguished by its novel integration of experimentally recorded feedback from the higher lateromedial (LM) visual area.

We started our analysis in L2/3 to simplify the analysis of LFPs generation mechanisms, and because previous computational work suggests that this layer provides a large contribution to extracellular potentials^53,67^ (Figure S1).

The L2/3 model was composed of a 400-μm radius inner cylinder of morphologically derived multi-compartmental neurons with somatic Hodgkin-Huxley dynamics and passive dendrites ^68^. This inner cylinder was surrounded by an annulus of point neurons whose purpose was to avoid boundary artifacts ^69^. L2/3 contained four distinct neuronal populations (Figure 1A, B), one of which was excitatory. To elaborate, the L2/3 network comprised a single model of excitatory cells, three models for the parvalbumin (PV) positive interneurons, four models for the somatostatin (Sst) positive interneurons, and eight models for the 5- hydroxytryptamine-expressing (Htr3a) interneurons (see Table 1 for populations numerosity). Overall, 85% of the neurons were excitatory and 15% were inhibitory.

**Figure 1.**
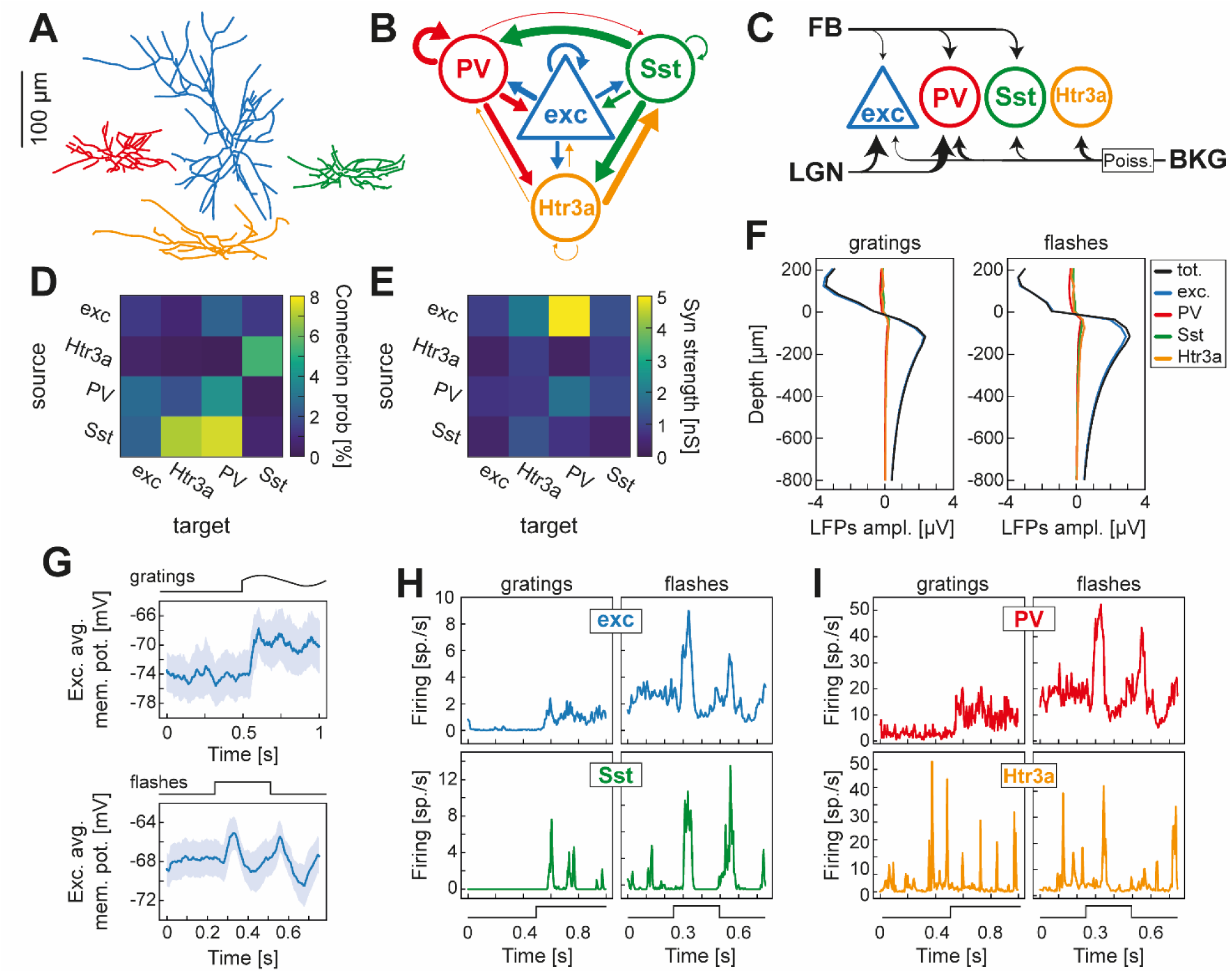
Features of mouse V1 layer 2/3 computational model. A. Representative morphological reconstructions of the four classes of L2/3 neurons: excitatory (blue, N=12689 cells, 1 model); parvalbumin (PV) positive interneurons (red, N=640 cells, 3 models); somatostatin (Sst) positive interneurons (green, N=464 cells, 4 models); Htr3a interneurons (orange, N=1107 cells, 8 models). B. Connectivity patterns of L2/3 network. The size of the arrows is proportional to the connection probability among neuronal classes. C. The L2/3 V1 model received three external stimuli: (i) feedback activity from lateromedial visual areas (FB); (ii) lateral geniculate nucleus afferents (LGN); (iii) background input. The spike trains driving the FB and LGN afferents were taken from publicly available experimental recordings, as described in (Rimehaug et al., 2023)^59^. Background input consisted instead of independent realizations of a homogeneous Poisson (‘Poiss.’) spike generator process with constant rate of 1000 sp./s. The width of the arrows contacting L2/3 neuronal cells are proportional to the synaptic strength of the three external stimuli. D. Average connection probabilities among L2/3 neuronal classes. E. Median synaptic strengths among L2/3 neuronal classes. F. Amplitudes of LFP signals generated in the morphological MC network across cortical depth when the network was presented with drifting gratings (left) and flashes of lights (right). LFP amplitudes were decomposed into the amplitude generated by transmembrane currents in all neurons (black, ‘tot.’ label), excitatory neurons (blue, ‘exc’), parvalbumin (red, ‘PV’), somatostatin (green, ‘Sst’), and Htr3a interneurons (orange, ‘Htr3a’). The amplitude was measured as the standard deviation of the LFP signals over the entire time course with the same sign as the mean of the signal at the corresponding cortical depth. The depth axis was centered around the average somatic position of L2/3 cells. G. (top) Average membrane potential of L2/3 excitatory cells in response to drifting gratings. The stimulus was composed of 500 ms of gray screen preceding 500 ms of sinusoidal drifting gratings. The gratings were characterized by a spatial frequency of 0.04 cycles per degree and by a temporal frequency of 2 Hz (see Methods). The temporal course of the stimulus is represented on top of the panel. Shaded region indicates inter quantile range of membrane potentials distribution of L2/3 excitatory cells. (bottom) Average membrane potential of L2/3 excitatory cells in response to full-field flashes. The flash stimuli trials consisted of 250 ms of gray screen, followed by 250 ms of white screen, returning to a gray screen for 250 ms (see Methods). The temporal course of the stimulus is represented on top of the panel. Shaded region indicates inter quantile range of the membrane potential distribution of L2/3 excitatory cells. H. Average instantaneous firing rate of L2/3 excitatory (top row, blue traces) and Sst interneurons (bottom row, green traces) in response to drifting gratings (left column) and flashes of light (right column) stimuli. The temporal course of the visual stimuli is schematically represented at the bottom of each column. I. Same as H, for parvalbumin (top row, red traces) and Htr3a (bottom row, orange traces) interneurons.

**Table 1.** Numerosity of populations in the V1 network model.

|  | N |
| --- | --- |
| L2/3: excitatory cells | 12689 |
| L2/3: PV interneurons | 640 |
| L2/3: Sst interneurons | 464 |
| L2/3: Htr3a interneurons | 1107 |
| LIF L2/3: excitatory cells | 43368 |
| LIF L2/3: PV interneurons | 2287 |
| LIF L2/3: Sst interneurons | 1656 |
| LIF L2/3: Htr3a interneurons | 3738 |
| L4: excitatory cells | 10254 |
| L4: PV interneurons | 963 |
| L4: Sst interneurons | 553 |
| L4: Htr3a interneurons | 270 |
| LIF L4: excitatory cells | 35507 |
| LIF L4: PV interneurons | 3498 |
| LIF L4: Sst interneurons | 1831 |
| LIF L4: Htr3a interneurons | 961 |
| L5: excitatory cells | 7569 |
| L5: PV interneurons | 613 |
| L5: Sst interneurons | 555 |
| L5: Htr3a interneurons | 117 |
| LIF L5: excitatory cells | 25989 |
| LIF L5: PV interneurons | 2263 |
| LIF L5: Sst interneurons | 1983 |
| LIF L5: Htr3a interneurons | 391 |

The network received external inputs from three different external sources (Figure 1C, see Methods): (i) experimentally recorded afferent activity from the lateral geniculate nucleus (LGN); (ii) experimentally recorded feedback from the lateromedial (LM) area of the visual cortex; and (iii) Poisson background spiking activity representing the (stimulus-independent) continuous influence of the rest of the brain on L2/3 V1. Recurrent connection probabilities (whose average values are reported in Figure 1D) were dependent on intersomatic distances as well as neurons’ functional preferences, such as cells’ direction tuning. Synaptic strengths (see Figure 1E for average values), on the other hand, followed an orientation-dependent like-to-like rule (i.e., cells preferring similar stimuli were preferentially connected^66^).

We simulated the network dynamics using two sensory stimuli: drifting gratings and full-field flashes. Flashes induced higher firing rates in every L2/3 neuronal family (except for Htr3a interneurons) (Figure 1I, J). This discrepancy was largest immediately after stimuli onsets and offsets. Flashes also induced a higher average membrane potential in excitatory neurons (Figure 1G, H).

This difference in neuronal response may be attributed to how the external inputs were modeled. For both stimuli, the FB inputs from the LM visual area were derived from experimentally recorded spike trains. However, the feedforward inputs from LGN were handled differently. In the case of flashes, the LGN spike trains were recorded experimentally in response to actual flashes of light. Conversely, for the drifting gratings, LGN inputs were modeled as in (Billeh et al., 2020)^66^ using the FilterNet module, which can generate LGN spike trains for a wide range of visual stimuli.

We found that, irrespective of stimulus, the amplitude of LFPs (obtained by simulating transmembrane currents using detailed multicompartmental neuron models) generated by L2/3 excitatory neurons were significantly larger than those generated by interneurons across the whole cortical depth (Figure 1F). In fact, the contribution to the overall LFPs from synaptic inputs onto every interneuron type (and their associated return currents) was minimal, consistent with what had been previously shown for stellate cells with symmetrically placed synapses^6,67^ (i.e., a so-called close-field arrangement^70^). Similarly, we observed that the LFPs generated by excitatory neurons in the deeper cortical layers also dominate those generated by interneurons (Figure S1), further emphasizing the minimal contribution of interneurons to the overall LFPs throughout the cortical depth. Accordingly, when we refer to the LFPs within a cortical layer, we are specifically referring to those generated from synaptic inputs onto the excitatory cells in that layer.

### Kernel-Based LFP estimation in detailed large-scale L2/3 network model

After simulating the LFPs generated by the highly biophysically detailed multicompartmental L2/3 network of the V1 model ^59,66^, we evaluated whether these signals could be effectively approximated using the computationally efficient kernel-based method introduced by (Hagen et al., 2022)^55^. This framework provides a computationally efficient method for estimating LFPs by convolving neuronal firing rates with a set of spatiotemporal kernels, which represent the averaged spike-triggered extracellular potential response between pre- and post-synaptic populations (see Methods for details).

In the original work by (Hagen et al., 2022) ^55^, this kernel-based method was tested on a minimalistic network consisting of only two types of neuronal populations (one excitatory and one inhibitory) subject to constant external synaptic input. The model also relied on several key simplifications, including fixed connection probabilities and uniform synaptic weights between connected populations. In contrast, the network examined in this study was vastly more intricate, encompassing 16 distinct neuronal sub-populations across six cortical layers, three types of time-varying external inputs, and a non-trivial, asymmetrical distribution of both connection probabilities and synaptic strengths between pairs of connected neurons. This added complexity reflects a more biologically realistic scenario, presenting a significant challenge for the kernel-based framework.

Before applying the kernel-based method, we computed two key network parameters: the average connection probability between neuronal populations (Figure 1D) and the median synaptic strength (Figure 1E). Given the significant skew in synaptic strength distributions (particularly among excitatory cells) we chose to use the median as a measure of central tendency, which more accurately represents the central behavior in these asymmetrical distributions (see Figure S2 for detailed distributions).

To estimate a set of LFP kernels for each pair of pre- and post-synaptic populations in L2/3, we followed the methodology described by (Hagen et al., 2022) ^55^ (see also Methods). As previously noted, we focused specifically on kernel pairs where L2/3 excitatory cells were the post-synaptic targets.

For each population connected to the L2/3 excitatory neurons, we calculated the corresponding set of LFP kernels. These kernels describe the average extracellular response in excitatory neurons to incoming synaptic activity from a given pre-synaptic population.

In addition to the structural parameters of the network (such as connection probabilities, synaptic delays, synaptic strengths, and the distribution of synaptic contacts along the morphology of target neurons), the kernel estimation process also require specifying the membrane potentials of the target populations, which affect the driving force of synaptic currents, as well as the firing rates of pre-synaptic populations, as they influence the effective leak conductance of post-synaptic neurons. Initially, we estimated LFP kernels using fixed values of both the membrane potentials of the target population and the presynaptic firing rates, consistent with the approach used by (Hagen et al., 2022) ^55^.

Although this approximation disregards the observed temporal fluctuations (Figure 1 G-J), the LFPs estimated by convolving the firing rates of each pre-synaptic population with its corresponding set of kernels provided a satisfactory approximation of the LFPs in response to flash stimuli (median R² = 0.82, Figure 2A). However, for visual gratings, the kernel-based estimation failed to capture the LFPs accurately (median R²=0.28, Figure 2B). We hypothesized that this limitation of the kernel-based approach arose from its core assumptions of constant membrane potentials and presynaptic firing rates. These factors directly influence synaptic currents and the passive properties of the post-synaptic membrane, which are known to fluctuate dynamically in response to naturalistic external inputs. To overcome this, we extended the kernel-based framework to account for these network non-stationarities.

**Figure 2.**
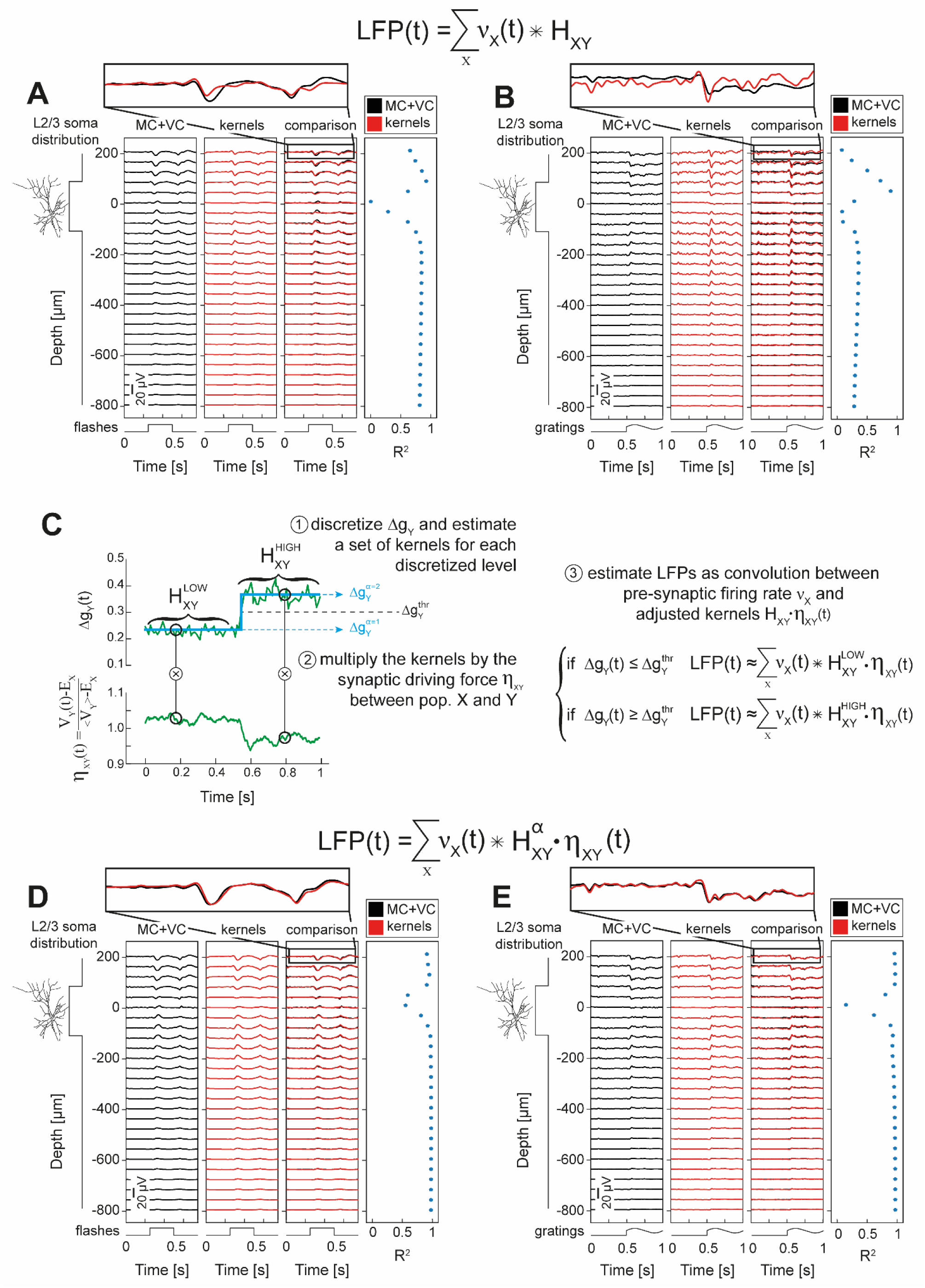
Mouse V1 layer 2/3 LFPs can be computed as the convolution between firing rates and dynamic spatiotemporal kernels. A. Depth-resolved trial-averaged L2/3 LFPs in response to full-field flashes of light. The temporal pattern of the visual stimulus is illustrated at the bottom of each panel. The LFPs were estimated using two methods: (i) biophysically detailed multi-compartmental (MC) neuronal network simulations combined with a forward model derived from volume conduction (VC) (black traces), and (ii) convolution of neuronal firing rates with a set of spatiotemporal kernels (red traces). The kernel-based estimation of LFPs employed a single set of kernels, assuming constant target membrane potential V_Y_(t) and presynaptic firing rates υ_X_(t). This estimation (see formula reported at the top of the panel) consisted of the sum of the convolutions between the presynaptic firing rates υ_X_(t) and the kernels HXY for the connection between presynaptic population X and target population Y. The consistency between the kernel-based and multicompartmental LFPs estimation was evaluated with R^2^ (right column). L2/3 somatic distribution along the cortical depth, graphically indicated on the left, was uniform within the [-105,105] µm range. The top inset provides a zoomed-in comparison between the LFPs estimated with MC simulations and kernels at the highest simulated recording electrode contact. B. Same as A, but in response to drifting gratings. C. The estimation of dynamic kernels can be broken down into three key steps: (1) The change in effective leak conductance, Δ_gX_(t), induced by presynaptic spiking activity, is discretized into N levels (e.g., N = 2 in this example). For each level, a corresponding set of kernels is estimated; (2) at each simulated time point, the selected kernels (based on the discrete level assumed by Δ_gX_(t)) are multiplied by the normalized synaptic driving force η_XY_(t)=(V_Y_-E_X_)/(<V_Y_>-E_X_) between the presynaptic population X and the postsynaptic population Y; (3) finally, LFPs are computed as the convolution between the presynaptic firing rate, υ_X_(t), and the product of the kernels H_XY_^α^ and the synaptic driving force η_XY_(t). The kernels are chosen dynamically over time, according to the discrete level α of Δg_X_(t). D. Same as A, but estimating LFPs using the procedure introduced in C (see the formula above the panel). E. Same as D, in response to drifting gratings.

### Dynamic kernels to enhance LFP estimation

We investigated how changes in two key factors affected LFP kernels: first, the membrane potential of the target population, and second, the firing rates of the presynaptic populations.

When calculating the kernels H_XY_ between a presynaptic population X and a postsynaptic population Y, the (assumed constant) mean membrane potential of the postsynaptic population <V_Y_> needs to be specified ^55^. This is because the postsynaptic currents are calculated from the synaptic driving force term <V_Y_>-E_X_, where E_X_ represent the reversal potential of the synapse between population X and Y. We found that the kernels scale linearly with changes in the synaptic driving force term (Figure S3A), indicating that fluctuations in V_Y_ can be accounted for without the need to re-estimate the kernels. Instead, we can simply re-scale the kernels (computed assuming constant <V_Y_>) using the time-variable postsynaptic membrane potential, expressed as:

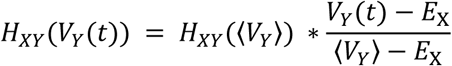

In contrast, fluctuations in presynaptic firing rates can be expected to introduce nonlinear effects on LFP kernels (Figure S3A). This occurs because presynaptic firing activity influences the effective leak conductance of the target neuron, thereby altering its passive membrane properties and ultimately leading to a nonlinear effect on kernel dynamics. More specifically, the change in effective leak conductance Δg_Y_(t) induced by presynaptic spiking activity f_X_(t) is described by the following equation (see Methods and ^55^):

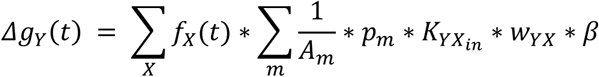

Where A_m_ is the area of compartment m, K_YXin_ is the per neuron indegree between presynaptic population X and postsynaptic population Y, w_YX_ is the synaptic weight, p_m_ is the per compartment connection probability between X and Y, and β is the integral of the sum of exponentials describing the synaptic conductance evolution.

The absence of a simple, direct relationship between kernel dynamics and presynaptic firing rate fluctuations implies that a different set of kernels would need to be recalculated at each time point, or at least for every value of Δg_Y_(t), which could undermine the computational efficiency of the kernel-based approach. To overcome this challenge, we propose discretizing the effective leak conductance Δg_Y_(t) into N discrete levels (see Methods). This approach allows LFP kernels to be computed for each distinct level, or for each time interval in which Δg_Y_(t) remains within the same discrete level, reducing the computational burden while still capturing key dynamics.

Thus, a time-variable version of the LFP kernel-based prediction can be expressed as:

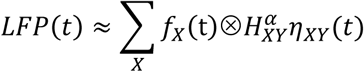

with 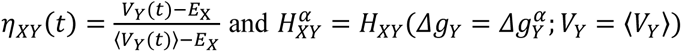 representing the set of kernels calculated for each discrete level α of Δg_Y_, assuming as target membrane potential its temporal average 〈𝑉_𝑌_〉.

This novel dynamic kernel estimation approach significantly improved the goodness-of-fit of LFP predictions for both drifting gratings (median R²=0.96, Figure 2D) and flashes of light (median R²=0.98, Figure 2E) with only N=2 discrete levels of Δg_X_(t).

Interestingly, we observed a notable drop in goodness-of-fit between LFPs estimated through kernel convolution and those simulated using multicompartmental models at positions near the midpoint of the somatic depth distribution (i.e., 0 on the depth axis in Figures 2D, 2E). This decline in R^2^ coincided with lower amplitude LFPs, and is likely attributable to the inversion point of the evoked transmembrane currents, leading to noisier, low-amplitude fluctuations in this region ^65^.

Overall, this time-varying kernel approach underscores the importance of accounting for network dynamics when exploring the mechanisms underlying extracellular potential generation, particularly when simulating variable-rate external inputs that induce highly non-stationary activity in neuronal networks, as is often the case in biological systems.

### LFPs in L2/3 mouse V1 model are dominated by synaptic feedback from higher visual areas

The kernel method allows the LFP to be represented as the sum of convolutions between the firing rates of individual neuronal subpopulations and the corresponding LFP kernels. This offers the possibility to discern the relative contribution of each neuronal family in shaping the overall LFP. Importantly, this can be achieved without the need to silence specific neuronal populations or modify the network structure in any way.

We applied the kernel method to disentangle the contributions of each L2/3 neuronal subpopulation to the collective LFPs in the mouse V1 L2/3 network model. In addition to the full LFP, we also decomposed the LFPs into: (i) the internal LFPs, generated by synaptic activity from within the L2/3; and the (ii) external LFPs, generated by synaptic inputs from external inputs.

For both stimuli, the kernels describing the effect of PV synaptic activity displayed the largest amplitude among internal populations (Figure 3A,G, and Figure S3B,C). Coupled with the fact that PV cells exhibited the highest firing rates among the V1 populations (Figure 1H,I), we found that the internal LFP variance was predominantly influenced by the activity of parvalbumin (PV) interneurons across the entire cortical depth (Figure 3C, I). In contrast, the contributions from the synaptic inputs of the other two families of inhibitory cells, Sst and Htr3a interneurons, and of excitatory cells were limited (Figure 3C, I). Consistently, the LFPs generated by PV interneurons explained nearly the entire variance of internal LFPs (Figure 3F, R^2^=0.99; Figure 3L, R^2^=1.0), while the remaining L2/3 populations accounted for a smaller proportion of the explained variance (Figure 3F, mean R^2^=0.07; Figure 3L, mean R^2^=0.20).

**Figure 3.**
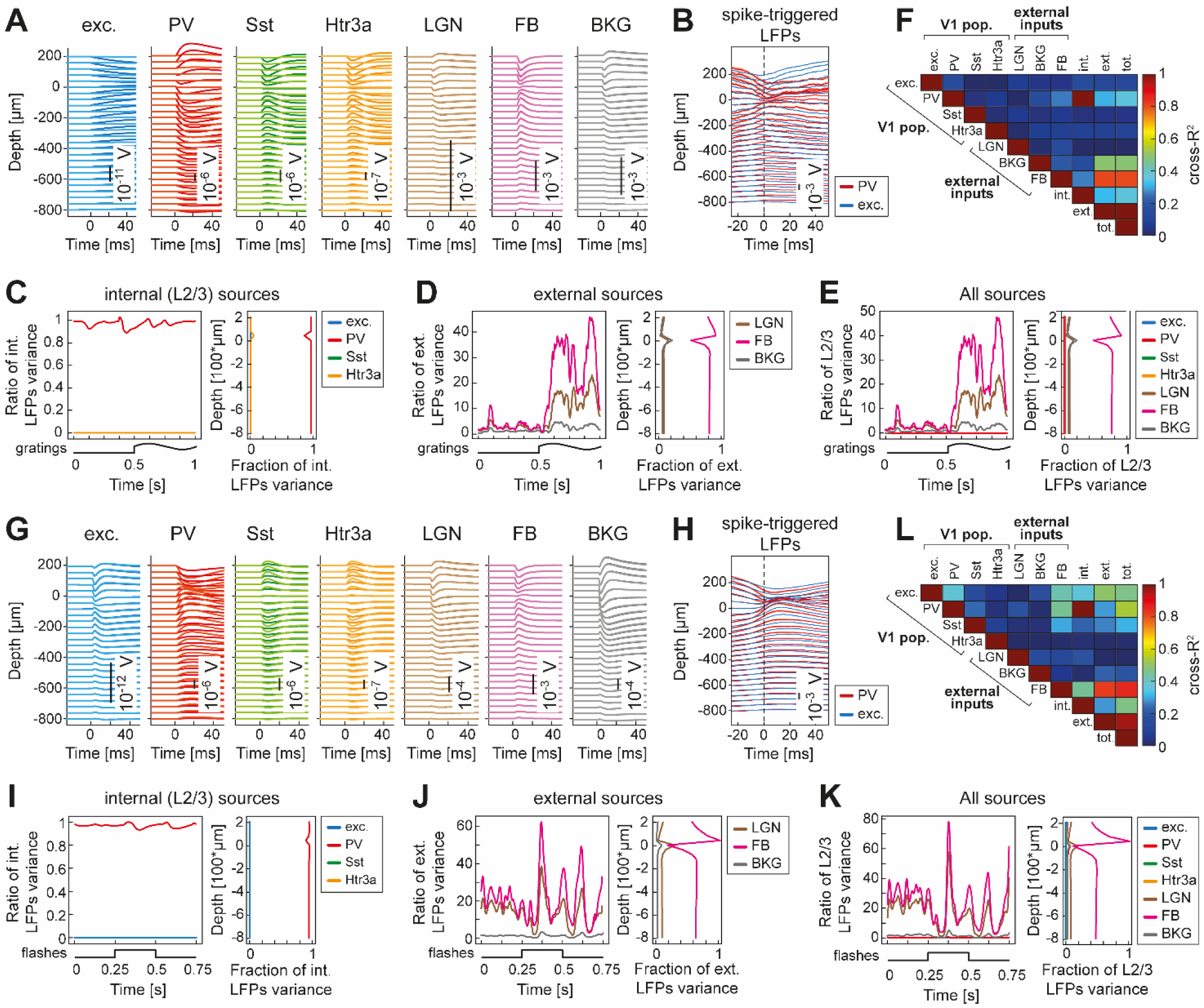
Contributions of multiple neuronal populations shape LFPs in the single-layer 2/3 model. A. Average spatiotemporal kernels for both discrete levels of 𝛥*g*_*X*_ (shown in different shades), used to compute LFPs in response to drifting gratings for each neuronal LFP source. The sources are (from left to right): excitatory cells (blue); PV (red), Sst (green), and Htr3a (orange) interneurons; LGN (brown), FB (violet), and BKG (grey). Kernels magnitudes are reported in the inset of each panel. B. Average LFPs triggered by spikes of L2/3 PV (red line) or L2/3 excitatory (red line) neurons. Shaded areas indicate the standard error of the mean. C. Variance ratios between internal LFPs (in response to drifting gratings) and the LFPs generated by the activity of excitatory cells (blue trace), PV (red trace), Sst (green trace), and Htr3a interneurons (orange trace). Internal LFPs are defined as the LFPs generated by neuronal populations within L2/3 of V1. (Right) Variance ratios between internal LFPs and L2/3 cortical neuronal populations across the cortical depth. Note that, because the variance of a sum includes additional covariance terms between components, the variance ratios do not sum to 1. Additionally, the variance ratios for Sst and Htr3a interneurons are approximately zero and are masked below the variance ratios of excitatory cells and PV interneurons. D. (left) Ratio of variance between external LFPs, in response to drifting gratings, and the LFPs generated by the inputs coming from LGN (brown trace), FB from LM (violet trace), and BKG (grey trace). (right) Ratio of variance between external LFPs and the external inputs across cortical depth. E. (left) Ratio of variance between the total LFPs, in response to drifting gratings, and the LFPs generated by the activity of every neuronal family in the model: excitatory cells (blue trace); PV interneurons (red trace); Sst interneurons (green trace); Htr3a interneurons (orange trace); LGN (brown); FB (violet); BKG (grey). (right) Ratio of variance between total LFPs and every neuronal family in the single-layer L2/3 model across cortical depth. F. R^2^ matrix between the overall LFPs and the LFP contributions generated by the firing rates of each neuronal population in the reduced single-layer V1 model. Please note that the labels: ‘int.’ indicate the sum of the LFPs generated by L2/3 neuronal populations; ‘ext.’ indicates the sum of the LFPs generated by external sources; ‘tot.’ implies the sum of ‘int.’ and ‘ext.’ LFPs. G-L. Same as A-F, in response to flashes of light.

We found the negligible contribution of Sst and Htr3a neurons to the LFPs could be attributed to several factors, all of which would need to be addressed to match the LFP amplitudes of PV neurons (Figure S4B,C): lower average firing rates (Figure 1I-J), weaker synaptic strength onto pyramidal cells (Figure 1B), and less spatially concentrated synapses (see Figure S4B,C and Methods). Unlike the internal LFP, the external LFP was not dominated by the synaptic activity coming from a single population. Nevertheless, the feedback (FB) activity from higher visual areas, that is, the lateromedial area (LM), emerged as the most significant contributor across the entire cortical depth for both stimuli, followed by the contributions coming from LGN and the background activity (BKG) (Figure 3D, J). This difference in the significance of these populations outside V1 became particularly evident during the presentation of drifting gratings (i.e., for t>500 ms in Figure 3D) and following the onset/offset of light flashes (i.e., for t>250 ms and t>500 ms in Figure 3J). Accordingly, the external LFPs variance was primarily explained by the feedback from LM inputs (Figure 3F, R^2^=0.84; Figure 3L, R^2^=0.85), followed by the background (Figure 3F, R^2^=0.49; Figure 3L, R^2^=0.22), and the LGN afferents (Figure 3F, R^2^=0.003; Figure 3L, R^2^=0.001).

Note that the variance of the external LFPs was lower than the sum of the variances of its components. Accordingly, the ratio between the variances of the external LFPs and the LFPs generated by the synaptic activity coming from each individual external populations exceeded 1. This arose because the LGN and BKG produced fluctuations with opposite signs to those from FB, as evidenced by the differing signs of their kernels (Figure 3A, G, and Figure S3B,C). This was due to synaptic positioning: LGN and BKG synapses were placed on the basal and apical dendrites of L2/3 pyramidal cells, while FB synapses only contacted the apical dendrites.

Overall, the total LFPs (i.e., the sum of internal and external LFPs) were primarily driven by neuronal populations outside V1, with internal populations playing only a marginal role (Figure 3E, K). The external inputs collectively accounted for nearly all the variance in the total LFPs (Figure 3F, R^2^ = 1.0; Figure 3L, R^2^ = 0.97), whereas the contribution from L2/3 neuronal populations explained a much smaller ratio of the variances (Figure 3F, R^2^ = 0.35; Figure 3L, R^2^ = 0.46) for both tested stimuli. Even the most influential internal source, the PV interneurons, explained relatively little variance (Figure 3F, R^2^ = 0.35; Figure 3L, R^2^ = 0.55). In contrast, the primary source of LFPs originated from feedback from the LM (Figure 3F, R^2^ = 0.84; Figure 3L, R^2^ = 0.89).

Interestingly, while other external inputs contributed significantly more to the total LFP variance than PV neurons (Figure 3E, K), the R^2^ between the total LFPs and the LFPs generated by PV interneurons (Figure 3F, R^2^ = 0.35; Figure 3L, R^2^ = 0.55) was higher than that of background inputs in response to flashes (Figure 3L, R^2^ = 0.20) and LGN inputs in response to both stimuli (Figure 3F, R^2^ = 0.003; Figure 3L, R^2^ = 0.001). This suggests that, while LGN and background inputs produced higher-amplitude LFPs, the LFPs generated by PV interneurons exhibited greater covariance with the total LFPs. This likely arose from the strong correlation between the firing rates of PV neurons and external inputs (Pearson’s correlation, ρ = 0.72, which increased to ρ = 0.76 at lag = 6 ms). However, the reverse did not hold true: while excitatory neurons exhibited a similar correlation with external stimuli (Pearson’s correlation, ρ = 0.73, increasing to ρ = 0.79 at a 9 ms lag), the LFPs associated with their synaptic activity showed negligible co-variation with the overall LFP.

This minimal contribution of local synaptic activity to the LFP might appear to contradict prior studies that emphasized the role of PV synaptic inhibition in shaping cortical LFPs ^53,63,71^. We suggest that the strong drive of PV neurons by external inputs may inflate their apparent contribution to the LFP. To explore this, we computed spike-triggered LFPs (st-LFPs), that is the average of short LFP segments centered around each spike time of PV and excitatory cells. We found that st-LFPs from both PV and excitatory neurons were three orders of magnitude larger than their respective kernels (Figure 3B, H; Figure S5), aligning more closely with the amplitudes of kernels associated with external inputs (Figure 3A, G). Further, in agreement with ^71^, we observed that st-LFPs from excitatory and PV cells exhibited the same polarity and similar amplitudes, unlike their respective kernels (Figure 3A, G vs. Figure 3B, H). Additionally, st-LFPs from PV cells slightly preceded those from excitatory cells, especially in response to flashes (Figure 3H). This timing difference likely reflects the longer transmission times in pyramidal cells, as also indicated by the correlation analysis above. Overall, these results imply that spike-triggered measures may misattribute LFP contributions from external inputs to PV neurons. In other words, st-LFPs recorded from PV cells can primarily reflect external synaptic input, while the postsynaptic LFPs generated by PV neurons are better captured by kernel-based estimates.

In summary, the LFPs in the mouse L2/3 network, as modeled here ^59,66^, are predominantly shaped by synaptic activity from populations external to V1, particularly feedback from the LM. Importantly, while this insight was easy to extract from the kernel-based LFP, it would have been much more difficult to extract from the original model, since the standard approach to model LFPs doesn’t provide easy access to the relative importance of different types of synaptic input in shaping LFP signals.

Of note, we compared the kernel method to a simplified LFP prediction approach based on linear combinations of synaptic currents from point-neuron network simulations, as previously proposed^63^. By designing a point-neuron network equivalent to the multicompartmental L2/3 network (Figure S6), we demonstrated that both approaches could accurately approximate internal LFPs (Figure S7: gratings median R^2^=0.90; flashes median R^2^=0.94). However, the synaptic current coefficients required optimization for each depth and network dynamics (Supplemental Text: “LFP proxies based on synaptic currents from point-neuron simulations”), which limits their generalizability. In contrast, the kernel method offered a robust and stimulus-independent solution for estimating LFPs from spiking activity in simplified neuronal models. Importantly, the kernels are deterministically derived from experimentally validated cell and network parameters, without any free parameters requiring fitting, further enhancing their general applicability and predictive power.

### Kernel-based approach captured L2/3 LFPs in a full-column model of mouse V1

Up to this point, we have focused on a cortical network limited to L2/3, excluding other layers present in the original model ^59,66^. To broaden our analysis, we applied the kernel-based method to estimate the LFPs generated by L2/3 pyramidal cells in the context of a full-column V1 model (see Methods).

This full-column model included six cortical layers, with L2/3 receiving additional inhibitory afferents from layers 1, 4, 5, and 6, and excitatory inputs from layers 4 and 5. As with the L2/3-only network, LFPs generated by inhibitory neurons were negligible, so we focused on L2/3 pyramidal cells as the primary contributors to the LFPs (Figure S1).

We derived spatiotemporal kernels using the same method as in Figure 2, discretizing the effective leak conductance 𝛥*g*_𝑌_(𝑡) into two levels (N=2) and convolving the firing rates with the corresponding kernels. This approach yielded accurate LFP estimates for L2/3 in the full-column model, both in response to contrast gratings (Figure 4A, R^2^=0.96) and flashes of light (Figure 4F, R^2^=0.98). As in the single-layer network, the goodness-of-fit decreased in the somatic depth region, likely due to noisy low-amplitude fluctuations.

**Figure 4.**
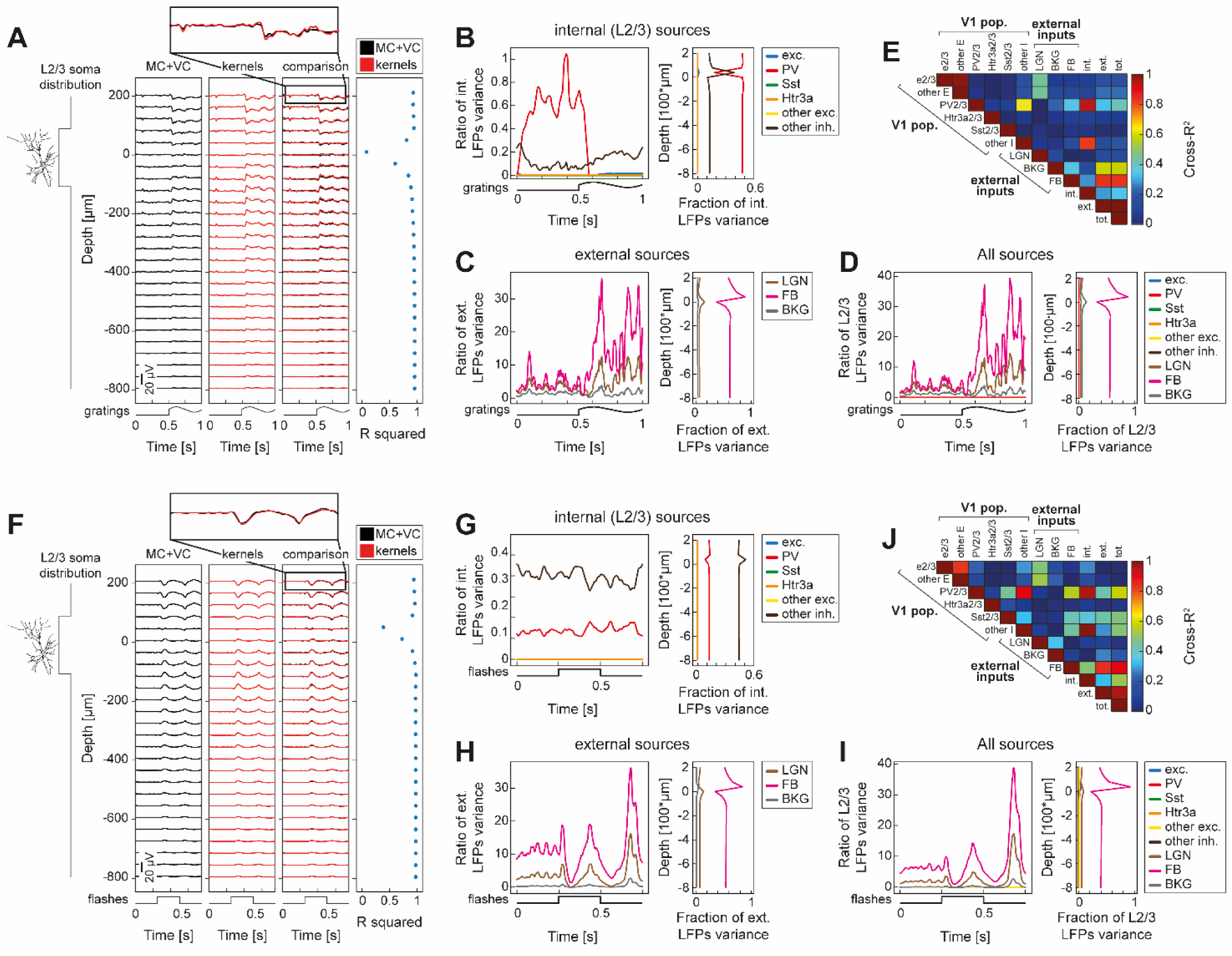
Layer 2/3 LFPs in a full-column model of mouse V1 can be computed as a convolution of firing rates and spatiotemporal kernels. A. Depth-resolved trial-averaged L2/3 LFPs in response to visual gratings in the mouse V1 full-column model. The temporal pattern of the visual stimulus is represented at the bottom of each panel. The LFPs were computed: (i) with biophysically detailed multi-compartmental (MC) neuronal network simulations combined with forward model derived from volume conduction (VC) (black traces); (ii) by the convolution of neuronal firing rates with spatiotemporal kernels (red traces). The consistency between the kernel-based and multicompartmental LFPs estimation was evaluated with R^2^ (right column). L2/3 somatic distribution along the cortical depth, graphically indicated on the left, was uniform within the [-105,105] µm range. The top inset provides a zoomed-in comparison between the LFPs estimated with MC+VC simulations and kernels at the highest simulated recording electrode contact. B. (left) Ratio of variance between L2/3 internal (i.e., neuronal sources within V1 full-column model) LFPs in the mouse V1 full-column model, in response to drifting gratings, and the LFPs generated by the activity of excitatory cells (blue trace); PV interneurons (red trace); Sst interneurons (green trace); Htr3a (orange trace) interneurons; inhibitory inputs coming from outside L2/3 (dark brown trace, labeled ‘other inh.’); excitatory inputs coming from outside L2/3 (yellow trace, labeled ‘other exc.’). (right) Ratio of variance across cortical depth between L2/3 internal LFPs and the cortical neuronal populations within the mouse V1 full-column model. C. (left) Ratio of variance between L2/3 external LFPs in the mouse V1 full-column model, in response to drifting gratings, and the LFPs generated by the inputs coming from LGN (brown trace), FB from LM (violet trace), and BKG (grey trace). (right) Ratio of variance across cortical depth between L2/3 external LFPs and the external inputs in the mouse V1 full-column model. D. (left) Ratio of variance between L2/3 total LFPs in the mouse V1 full-column model, in response to drifting gratings, and the LFPs generated by the activity of: excitatory cells (blue trace); PV interneurons (red trace); Sst interneurons (green trace); Htr3a interneurons (orange trace); inhibitory inputs coming from outside L2/3 (dark brown trace, labeled ‘other inh.’); excitatory inputs coming from outside L2/3 (yellow trace, labeled ‘other exc.’); LGN (brown); FB (violet); BKG (grey). (right) Ratio of variance across cortical depth between L2/3 LFPs and every neuronal family in the mouse V1 full- column model. E. R^2^ matrix between the overall L2/3 LFPs and the LFP contributions generated by the firing rates of each neuronal population in the mouse full-column V1 model. Please note that the labels: ‘int.’ indicate the sum of the LFPs generated by the activity of every V1 neuronal family; ‘ext.’ indicates the sum of the LFPs generated by the activity of the three external inputs; ‘other I’ indicate the LFPs generated by the activity of inhibitory inputs coming from outside L2/3; ‘other E’ indicates the LFPs generated by the activity of excitatory inputs coming from outside L2/3; ‘tot.’ indicates the sum of the LFPs ‘int.’ and ‘ext.’. F-J. Same as A-E, in response to flashes of lights.

We used the kernel method to disentangle the contributions from different neuronal families. The L2/3 internal LFPs (generated by synaptic activity from within V1) were mainly influenced by PV interneurons and inhibitory inputs from other layers (Figure 4B, G). PV inputs alone accounted for most of the internal LFP variance (Figure 4E, R^2^=0.94; Figure 4J, R^2^=0.96), while excitatory (Figure 4E, R^2^=0.07; Figure 4J, R^2^=0.27) and non-PV inhibitory activity (Figure 4E, R^2^=0.04; Figure 4J, R^2^=0.24) had negligible effects.

LFPs generated by external sources, particularly feedback from LM, dominated the L2/3 LFPs in the full-column model (Figure 4E, R^2^=0.87; Figure 4J, R^2^=0.88). Background input (Figure 4E, R^2^=0.60; Figure 4J, R^2^=0.37) and thalamic afferents (Figure 4E, R^2^=0.09; Figure 4J, R^2^=0.12) had smaller but notable contributions.

Overall, external inputs accounted for almost the entire L2/3 LFP variance in the full-column model (Figure 4E, R^2^=1.0 for gratings; Figure 4J, R^2^=0.96 for flashes, and Figure S8), while V1 synaptic activity explained less variance (Figure 4E: R^2^=0.34 for gratings, Figure 4J: R^2^=0.52 for flashes, and Figure S8). Feedback from higher visual areas, particularly LM, was the main driver of L2/3 LFPs, with only PV interneurons within L2/3 providing significant internal contributions (Figure 4E, R^2^=0.39; Figure 4J, R^2^=0.60, and Figure S8).

In summary, our version of the kernel method accurately estimated L2/3 LFPs within the full-column V1 model. Consistent with findings from single-layer models, we observed that L2/3 LFPs were predominantly driven by external inputs, particularly feedback from the LM area, while internal V1 activity played a minimal role.

### Kernel-based LFPs captured L4 LFPs in a full-column model of mouse V1

After establishing the suitability of the kernel-based L2/3 LFP estimation (both in the single-layer and the full-column model), we turned our attention to adjacent cortical layers, first the underlying cortical layer, L4.

In the full-column V1 model, L4 was akin to L2/3, adhering to the same rules governing connectivity and synaptic strength allocation. However, L4 exhibited higher heterogeneity in the composition of its neuronal populations. L4 was composed of 4 families of excitatory cells, 2 families of PV and Sst interneurons, and one for the Htr3a cells (see Table 1 for populations numerosity). As for L2/3, the ratio between excitatory and inhibitory neurons was (85:15).

A notable difference to L2/3 was that L4 did not receive any feedback inputs from higher visual areas, which were identified as the main contributor to L2/3 LFPs (Figure 3,5). Thus, L4 external inputs were solely composed of thalamic afferents and the background activity.

**Figure 5.**
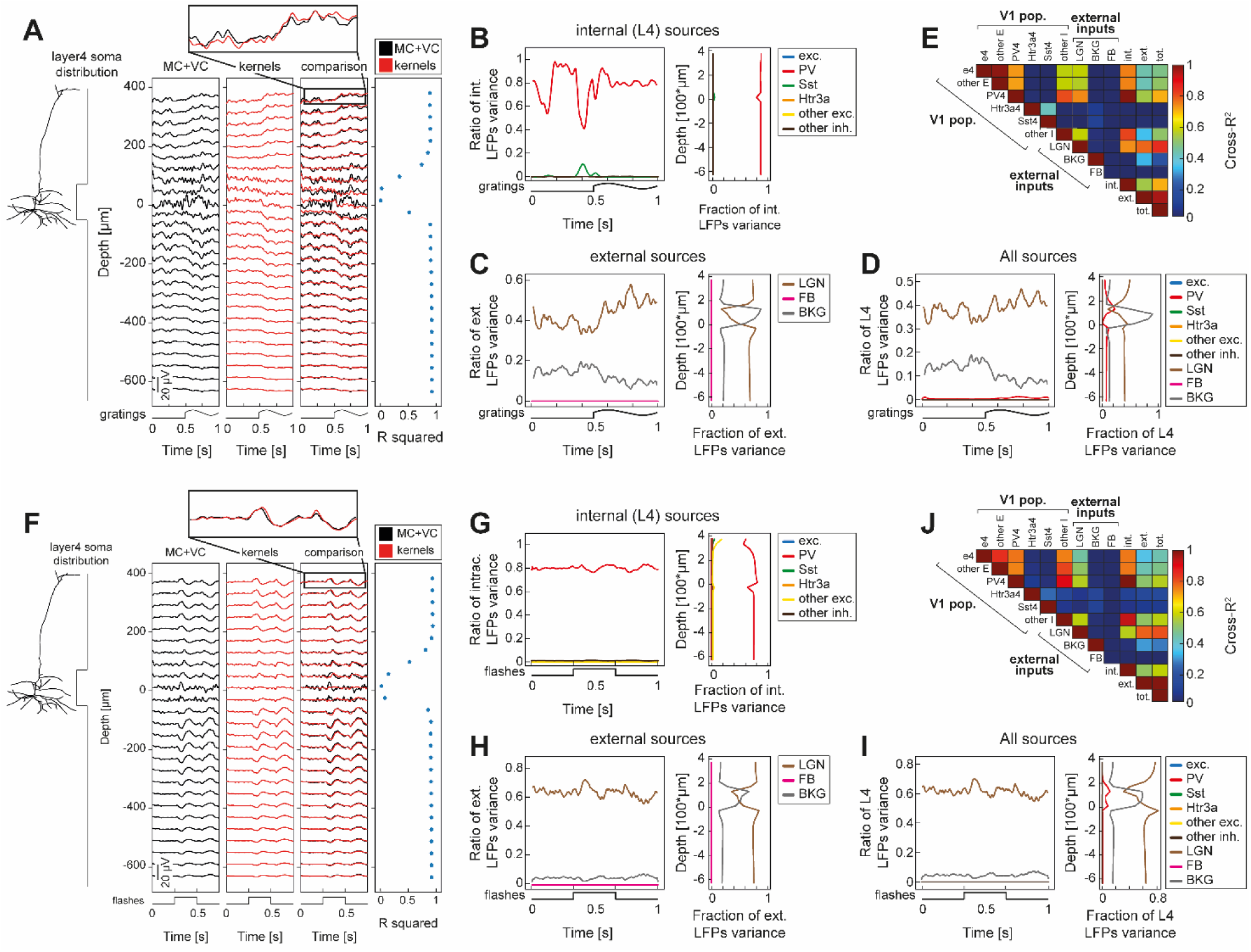
Layer 4 LFPs in a full-column model of mouse V1 can be computed as a convolution of firing rates and spatiotemporal kernels. A. Depth-resolved trial-averaged L4 LFPs in response to visual gratings in the mouse V1 full-column model. The temporal pattern of the visual stimulus is represented at the bottom of each panel. The LFPs were computed: (i) with biophysically detailed multi-compartmental (MC) neuronal network simulations combined with forward model derived from volume conduction (VC) (black traces); (ii) by the convolution of neuronal firing rates with spatiotemporal kernels (red traces). The consistency between the kernel-based and multicompartmental LFPs estimation was evaluated with R^2^ (right column). L4 neurons somatic distribution, graphically indicated on the left, was uniform within the [-60,60] µm range. The top inset provides a zoomed-in comparison between the LFPs estimated with MC+VC simulations and kernels at the highest simulated recording electrode contact. B. (left) Ratio of variance between L4 internal (i.e., neuronal sources within V1 full-column model) LFPs in the mouse V1 full-column model, in response to drifting gratings, and the LFPs generated by the activity of excitatory cells (blue trace); PV interneurons (red trace); Sst interneurons (green trace); Htr3a (orange trace) interneurons; inhibitory inputs coming from outside L4 (dark brown trace, labeled ‘other inh.’); excitatory inputs coming from outside L4 (yellow trace, labeled ‘other exc.’). (right) Ratio of variance across cortical depth between L4 internal LFPs (i.e., the LFPs generated by V1 neuronal populations) and the cortical neuronal populations within the mouse V1 full-column model. C. (left) Ratio of variance between L4 external LFPs in the mouse V1 full-column model, in response to drifting gratings, and the LFPs generated by the inputs coming from LGN (brown trace), FB from LM (violet trace), and BKG (grey trace). (right) Ratio of variance across cortical depth between L4 external LFPs and the external inputs to the mouse V1 full-column model. D. (left) Ratio of variance between L4 total LFPs in the mouse V1 full-column model, in response to drifting gratings, and the LFPs generated by the activity of: excitatory cells (blue trace); PV interneurons (red trace); Sst interneurons (green trace); Htr3a interneurons (orange trace); inhibitory inputs coming from outside L4 (dark brown trace, labeled ‘other inh.’); excitatory inputs coming from outside L4 (yellow trace, labeled ‘other exc.’); LGN (brown); FB (violet); BKG (grey). (right) Ratio of variance across cortical depth between L4 total LFPs and the neuronal populations of the mouse V1 full-column model. E. R^2^ matrix between the overall L4 LFPs and the LFP contributions generated by the firing rates of each neuronal population in the mouse full-column V1 model. Please note that the labels: ‘int.’ indicate the sum of the LFPs generated by the activity of every V1 neuronal family; ‘ext.’ indicates the sum of the LFPs generated by the activity of the three external stimuli; ‘other I’ indicate the LFPs generated by the activity of inhibitory inputs coming from outside L4; ‘other E’ indicates the LFPs generated by the activity of excitatory inputs coming from outside L4; ‘tot.’ indicates the sum of the LFPs ‘int.’ and ‘ext.’. F-J. Same as A-E, in response to flashes of lights.

Similarly to in L2/3, the L4 LFPs were mostly generated by the transmembrane currents of excitatory cells (Figure S1B). Consequently, we defined ‘L4 LFPs’ as the LFPs generated across the entire cortical depth by transmembrane currents in excitatory cells within layer 4. Accordingly, when estimating LFP kernels, we considered only L4 excitatory neurons as target populations.

We adopted the same dynamic estimation approach as described for the upper visual layers (see above and Methods) with N=2 discrete levels.

The kernel method accurately estimated L4 LFPs, both when presented with contrast gratings (Figure 5A, R^2^=0.90) and with flashes of lights (Figure 5F, R^2^=0.92). Similar to the situation for L2/3 LFPs, the goodness-of-fit of kernel-based LFPs estimation decreased at the cortical depth corresponding to the position of L4 cell somas (Figure 5A, F).

Internal L4 LFPs (i.e., the LFPs generated by synaptic activity coming from V1 neuronal populations) were predominantly shaped by L4 PV interneuron activity, exhibiting significant influence during both drifting gratings (Figure 5B) and flashes of light (Figure 5G). In fact, PV activity in this layer was the primary determinant of LFPs patterns (Figure 5E, R^2^=1.0; Figure 5J, R^2^=1.0).

LFPs generated by external synaptic inputs in L4 (Figure 5C, H) were primarily accounted for by LGN afferents (Figure 5E, R^2^ = 0.80; Figure 5J, R^2^ = 0.81), except for a region close to L4 cell somas, where external LFPs were predominantly shaped by background activity (Figure 5C, H). However, across the entire cortical depth and throughout the whole simulations, BKG activity only marginally influenced external LFPs (Figure 5E, R^2^ = 0.31; Figure 5J, R^2^ = 0.29).

Overall, the total L4 LFPs (i.e., the sum of internal and external LFPs) were primarily influenced by external synaptic inputs (Figure 5D, I), serving as the primary determinants accounting for a large fraction of L4 LFPs explained variance (Figure 5E, R^2^ = 0.96; Figure 5J, R^2^= 0.99). Thalamic afferents were the principal source of L4 LFPs (Figure 5E, R^2^= 0.88; Figure 5J, R^2^= 0.83). Similar to what was observed for L2/3, the sum of LFPs generated by every cortical neuronal family (Figure 5E, R^2^ = 0.72; Figure 5J, R^2^ = 0.58) was inferior to that explained by the LFPs induced by LGN alone. Interestingly, however, L4 PV interneurons represented the second-most important LFP source, explaining R^2^ = 0.72 (Figure 5E) in response to drifting gratings and R^2^ = 0.60 (Figure 5J) in response to full-field flashes of lights.

In summary, kernel-based LFP estimation proved accurate also in layer 4. While L4 LFPs, like those in L2/3, primarily reflected external synaptic inputs, they were predominantly shaped by excitatory synaptic activity from thalamic afferents (LGN), unlike in L2/3. Notably, PV interneurons in L4 accounted for a small fraction of the total L4 LFP variance (Figure 5D, I) while still maintaining the second-highest R-squared value (Figure 5E, J). This indicates that although the LFP is driven by external synaptic inputs, the contribution from PV interneurons closely covaries with the overall LFP.

### Layer 5 asymmetrical distribution of firing rates precludes accurate kernel-based LFPs estimation

We investigated kernel-based LFP estimation within L5 of the V1 model. As in layers 2/3 and 4, the transmembrane currents of pyramidal cells were the largest contributors to the overall LFPs (Figure S1C). Given this, we focused specifically on the LFPs generated by L5 excitatory cells, particularly the largest subfamily identified by the Rbp4 Cre-line, which almost entirely dominated the LFPs in L5. This neuronal population comprised 10 distinct subfamilies, each characterized by its unique electrophysiological model. The largest subfamily (referred to as subfamily 1 in Figure 6) encompassed 2350 cells, whereas the remaining nine subfamilies (referred to as subfamilies 2-10 in Figure 6) collectively counted 3660 cells.

**Figure 6.**
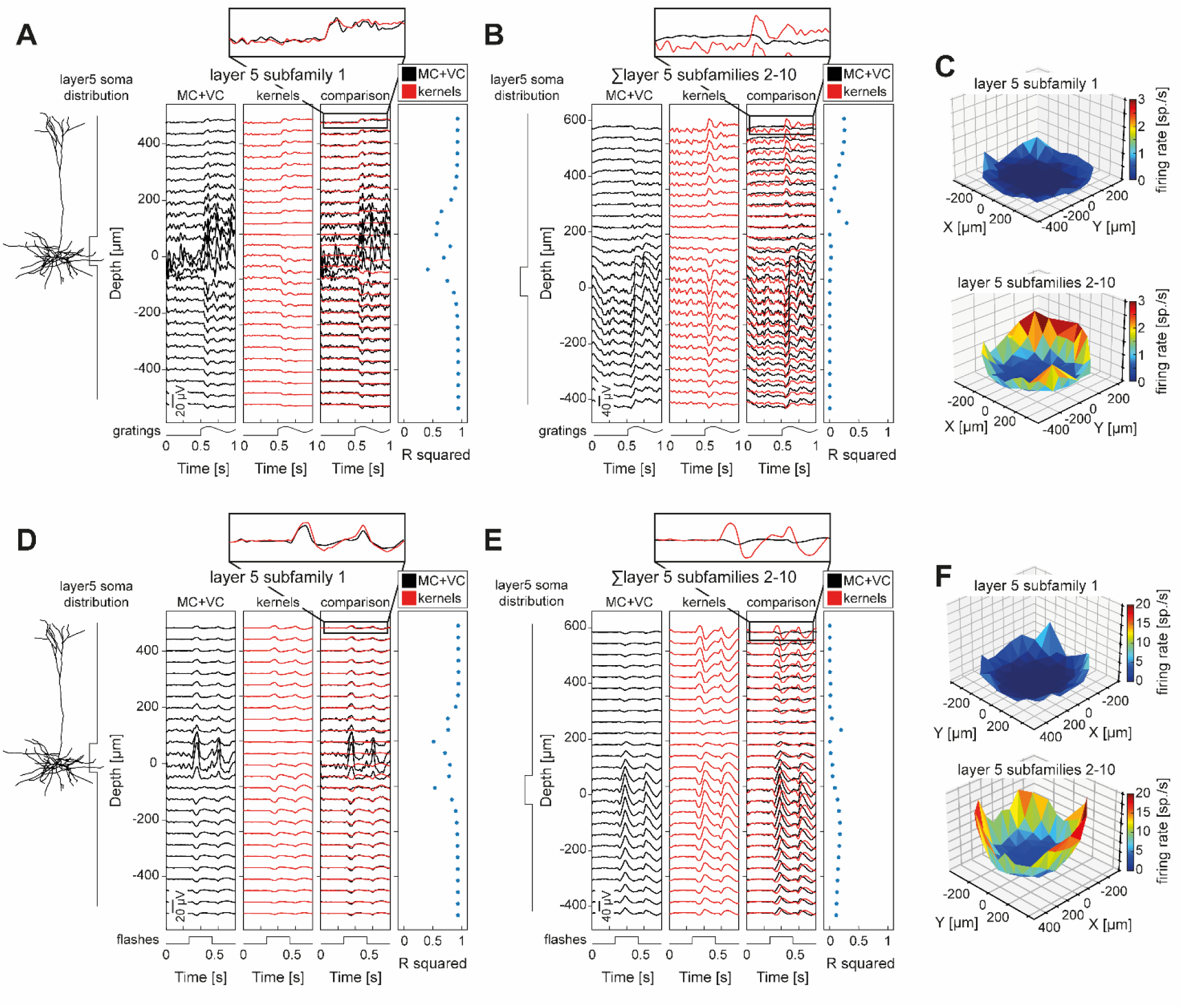
Layer 5 asymmetrical distribution of firing rates limits the effectiveness of the kernel-based estimation of LFPs. A. Depth-resolved trial-averaged LFPs in response to visual gratings in the mouse V1 full-column model. The LFPs were generated by the largest subfamilies of L5 excitatory cells, named ‘layer 5 subfamily 1’ in the panel title. The temporal traces of the visual stimulus are represented at the bottom of each panel. The LFPs were computed: (i) with biophysically detailed multi-compartmental (MC) neuronal network simulations combined with forward model derived from volume conduction (VC) (black traces); (ii) by the convolution of neuronal firing rates with spatiotemporal kernels (red traces). The consistency between the kernel-based and multicompartmental LFPs estimation was evaluated with R^2^ (right column). The somatic distribution of the largest subfamily of L5 excitatory cells, graphically indicated on the left, was uniform within the [-40,40] µm range. The top inset provides a zoomed-in comparison between the LFPs estimated with MC+VC simulations and kernels at the highest simulated recording electrode contact. B. Same as A, but the LFPs were the sum of the other nine L5 excitatory cells subfamilies, hence excluding L5 excitatory subfamily 1. The somatic distribution of these nine L5 neuronal subfamilies, graphically indicated on the left, was uniform within the [-70,70] µm range. C. Distribution of firing rate within L5 orthogonal plane of L5 excitatory subfamily 1 (top) and of the other nine L5 excitatory subfamilies (bottom). D-F. Same as A-C, in response to flashes of lights.

For each of these subfamilies, we estimated a distinct set of kernels, treating each subfamily as the target population, following the same approach used for L2/3 and L4 excitatory cells.

We observed that the set of kernels designed for ’subfamily 1’ as the target population accurately captured LFPs in response to both drifting gratings (Figure 6A, median R^2^ = 0.93) and flashes of light (Figure 6D, median R^2^ = 0.92), except near the depth of the layer 5 somata. In contrast, the goodness-of-fit for the remaining nine L5 pyramidal subfamilies was significantly lower (Figure 6B, median R^2^ = 0.004; Figure 6E, median R^2^ = 0.07) This inability in kernel-based LFP estimation might arise from the observation that the LFPs for these nine L5 subfamilies, as estimated through multicompartmental network simulations, displayed an unusual imbalance between negative and positive extracellular deflections across cortical depth (illustrated by the black traces in Figure 6B, E). This lack of balance was something particular to these subfamilies, as it was not displayed by the LFPs in L2/3 (depicted in Figure 2, Figure 4), L4 (illustrated in Figure 5A, F), or ’subfamily 1’ of L5 (shown in Figure 6A, D). This depth asymmetry in the LFP is a feature that the kernel-based LFPs estimation struggles to replicate, as evidenced by the unsatisfactory fit of kernel-based LFPs for L5 excitatory cells in the nine L5 “subfamilies 2-10” (Figure 6B, E). The LFPs estimated for these subfamilies with the kernel method (red traces in Figure 6B, D) were indeed characterized by a balanced depth profile, which evidently could not match the unbalanced pattern observed with multicompartmental network simulations. Importantly, an underlying assumption of the kernel method is that the cells of the postsynaptic population have uniform properties, are uniformly distributed within the plane of a cylinder, and receive the same kind of synaptic input. This leads to the transmembrane currents of the postsynaptic population to be uniformly distributed within the plane of a cylinder, while having a balanced depth profile due to current conservation ^2^. This, in turn, produces symmetrical negative and positive extracellular deflections across cortical depth. An imbalanced depth profile in the extracellular space, as observed in the LFPs produced by these L5 subfamilies, therefore suggests a significant departure from the assumptions underlying the kernel method adopted here.

The imbalance feature of these extracellular potentials likely originated in the asymmetrical distribution of excitatory cells firing rate across the L5 orthogonal plane. "Subfamily 1" (which is the only one for which kernel-based LFPs estimation proved effective) exhibited an approximately uniform distribution of mean firing rates across the x-y L5 plane for both stimuli (Figure 6C top, Figure 6F top). Specifically, only a minority of cells (16.37% in response to gratings and 34.43% in response to flashes) located more than 50 µm from the center of the L5 orthogonal plane displayed a firing rate twice as high as the average firing rate of those within a 50 µm distance. These fractions were comparable to those observed in L2/3 (24.47% in response to gratings and 10.45% in response to flashes) and L4 (17.19% in response to gratings and 19.57% in response to flashes). In contrast, the deviation from uniformity of firing rates across the L5 x-y plane was higher for the other nine L5 subfamilies (Figure 6C bottom, Figure 6F bottom). In these neuronal families, approximately one out of two cells closer to the visual column edges exhibited a firing rate twice as high as the average firing rate in a 50 µm inner cylinder (47.19% in response to gratings and 53.64% in response to flashes).

In turn, this lack of planar uniformity of firing rates of these L5 pyramidal subfamilies likely stem from an over-excitation feed coming from the external annulus of point neuron surrounding the central core of multicompartmental neurons (see Methods and (Billeh et al., 2020)^66^ for network architecture).

Overall, the kernel method utilized in this study was found to be ineffective in estimating LFPs when significant asymmetries are present orthogonal to the main axis along which neuronal morphologies are aligned.

Finally, we focused on subfamily 1, where the kernel-based LFP predictions were highly accurate, and analyzed the contributions of different presynaptic populations to the L5 LFPs (Figure S9). Consistent with our findings in L4, L5 LFPs were predominantly driven by external synaptic inputs (R^2^ = 0.95 for gratings and R^2^ = 0.93 for flashes), with thalamic afferents playing a major role. Among V1 populations, inhibitory inputs from L5 PV interneurons, as well as inhibition coming from other layers, were the most significant contributors to L5 LFPs. In contrast, excitatory synapses and inhibitory inputs from Sst and Htr3a interneurons had minimal to no impact.

## Discussion

The main outcome of our work was the demonstration of the feasibility of efficiently estimating LFPs using a kernel-based method in a large-scale, biophysically detailed model of the mouse V1. To account for network non-stationarities, we extended the original method by (Hagen et al., 2022)^55^ to make the kernels dynamic, allowing them to adapt in real time to changes in presynaptic firing rates and postsynaptic membrane potentials. This enhancement enabled the kernel method to accurately capture the complex, time-varying mechanisms driving LFP generation.

A second key point was the demonstration of how the kernel method can aid LFP analysis, which helped us reveal the minimal contribution of V1 populations to the overall LFP signal in the Allen V1 model. Instead, we found that external synaptic inputs dominated: L2/3 LFPs were primarily driven by feedback from higher visual areas, while L4 LFPs were shaped predominantly by thalamic inputs.

### Kernel-based scheme for predicting LFPs

#### LFPs can be efficiently estimated in a large-scale biophysically detailed computational model

One of the key advantages of the kernel-based framework is its ability to account for a high level of biological detail within a computationally efficient and straightforward structure. Unlike traditional multicompartmental network simulations, the kernel method estimates LFPs by convolving spatiotemporal kernels with neuronal firing rates. Notably, this approach decouples the estimation of extracellular potentials from the simulation of underlying neuronal dynamics. Consequently, neuronal activity can be modeled using simplified architectures, such as point-neuron or rate-based models, or even directly estimated from experimentally recorded spike trains, further reducing the computational burden associated with LFP estimation.

In this study, we applied the kernel-based method to estimate LFPs by convolving population firing rates with spatiotemporal kernels. This approach builds on a previous work ^55^, who first validated the kernel-based framework in a simplified network model consisting of single excitatory and inhibitory populations with steady external input. We extended this method to a more detailed and biologically realistic model of mouse V1, developed by the Allen Institute ^66^ and further refined by (Rimehaug et al., 2023)^59^. This large-scale V1 model comprises over 50,000 multicompartmental neurons distributed across six cortical layers and includes 17 distinct cell types, including multiple inhibitory neurons, whose contributions to cortical LFP generation had not been fully explored. With its unprecedented biological realism, the V1 model reproduces a wide range of experimental observations ^59,66^, making it an ideal testbed for validating the kernel-based LFP estimation approach.

A significant finding of our study is that the kernel-based method provided highly accurate LFP estimates in this complex, biophysically detailed model. A crucial insight from our work is that accounting for dynamic fluctuations in membrane potentials and firing rates is necessary for maintaining prediction accuracy. To address this, we developed a novel parametric extension of the kernel method, wherein passive membrane conductance changes in postsynaptic neurons, induced by ongoing synaptic inputs, were discretized into distinct levels. This discretization minimizes computational complexity while preserving the accuracy of kernel-derived LFPs.

A key contribution of this study is the substantial reduction in computational demands compared to traditional multicompartmental simulations. For instance, estimating LFPs via multicompartmental simulations in (Billeh et al., 2020)^66^ required approximately 90 minutes on 384 CPU cores to simulate 1 second of data. In contrast, we computed LFP kernels on a desktop computer in less than 10 minutes on 4 CPU cores. This drastic reduction in computation time highlights the efficiency and scalability of our method, making it highly advantageous for researchers. The efficiency gains extend even further, as the kernel method can produce LFP estimates when firing dynamics are simulated through simplified networks of point leaky-integrate-and-fire neurons or rate-based models.

Overall, the ability to drastically reduce computation time while maintaining accuracy holds important implications for the usability of detailed V1 models in the neuroscience community. This enhanced computational efficiency facilitates larger-scale and more flexible simulations of cortical dynamics, potentially advancing our understanding of brain function.

#### Comparison with other simplified approaches to estimate LFPs

As previously noted, the kernel method enables LFP estimation by convolving deterministic spatiotemporal kernels with neuronal firing rates, independent of how these rates are estimated or simulated. This allowed us to simulate the firing rates of the single-layer L2/3 V1 model using a point neuron network that functionally mirrored the multicompartmental model. By convolving the LFP kernels with these firing rates, we generated L2/3 LFP estimates.

Building on the work of ^63^, we demonstrated (Supplemental Text: “LFP proxies based on synaptic currents from point-neuron simulations”) that a linear combination of synaptic currents extracted from point neuron simulations could effectively predict the L2/3 LFPs obtained through kernel convolution (Supplemental Text: “LFP proxies based on synaptic currents from point-neuron simulations” and Figures S7, S10). This works because synaptic currents result from the convolution of presynaptic firing rates with synaptic conductance dynamic, which is typically expressed as sums of exponential functions (Mazzoni et al. 2008). Thus, the kernel method and the linear combination of synaptic currents share a conceptual basis: both approximate LFPs by convolving neuronal firing rates with biophysically derived kernels.

While our findings validate and extend the proxy method proposed by (Mazzoni et al., 2015)^63^ to networks containing multiple inhibitory neuron populations, they also highlight important limitations. Notably, we found that neglecting temporal non-stationarities reduced accuracy in LFP estimation. The need for dynamically adjusted linear combination coefficients underscores the significance of temporal fluctuations in neuronal activity, complicating straightforward applications of the proxy approach in scenarios where sensory stimuli induce substantial non-stationarities. This complexity raises questions about the reliability of using fixed coefficients of synaptic currents in dynamic environments.

In addition to these temporal challenges, we identified two further drawbacks of using LFP proxies based on linear combinations of synaptic currents. First, changes in sensory input can significantly alter network dynamics, necessitating recalibration of the coefficients for each condition. This need for recalibration limits the generalizability of the proxy model across different stimuli or experimental contexts.

Second, we highlight the crucial influence of neuronal morphology and synaptic placement on LFP patterns. The kernel method inherently accounts for these anatomical factors by integrating both neuronal structure and synaptic location into its predictions. In contrast, LFP proxies that rely on linear combinations of synaptic currents necessitate the engineering of new coefficients for every alteration in morphological parameters. This requirement restricts the adaptability of such proxy methods across different brain regions or networks with distinct architectures.

In conclusion, while the linear combination approach provides valuable insights, its practical applicability across diverse conditions and brain regions is limited due to its reliance on network-specific coefficients and structural parameters. The kernel method, on the other hand, offers a more flexible and robust framework for LFP estimation, accommodating a wider range of dynamic and anatomical influences, thereby enhancing its utility in neuroscience research.

### Results of *in silico* experiments on mouse V1 LFPs

The kernel approach allows for the decomposition of LFPs into contributions from individual neuronal populations, enhancing the interpretative power of LFP estimation. Leveraging this capacity, our investigation revealed that mouse V1 LFPs, as modelled here, are primarily driven by excitatory external inputs, notably feedback from higher-order visual areas in upper layers (Figure 3,5) and thalamic afferents in L4-5 (Figure 5, Figure S9). In contrast, local synaptic inputs from V1 populations contributed minimally to the overall LFP, although R^2^ analysis demonstrated strong covariance between the LFP generated by PV synaptic activity and the total LFP.

These findings align with the prevailing view that LFPs primarily reflect inputs to a neural network rather than its intrinsic activity ^1,3,72,73^, as corroborated by recent studies specifically focused on mouse V1 ^23,44,74,75^. However, our work advances this understanding by not only reaffirming that LFPs are driven by external inputs but also identifying their primary sources. Specifically, we found that the dominant contributions to V1 LFPs originated from brain regions outside V1 itself, such as higher-order visual areas and thalamic afferents, indicating that while V1 generates the recorded LFPs, the driving force behind these signals comes from external brain structures.

Our results thus reinforce the notion of complementarity in extracellular potentials, where V1 LFPs capture synchronous synaptic inputs from distant brain regions, while multi-unit and single-unit activities reflect locally generated action potentials. This complementary relationship is corroborated by multiple studies that have shown the synergistic contributions of these signals to the decoding of neuronal dynamics ^18,19,76,77^. Given the non-redundant information that LFPs and spikes provide about cortical dynamics, brain-machine interfaces could benefit from incorporating LFP signals, potentially enhancing neuronal decoding performance ^14,78^.

Multiple reports ^63,71,77,79,80^ have emphasized the role of inhibitory synaptic inputs, particularly from PV interneurons, in shaping LFP patterns. At first glance, this might seem at odds with our findings, which show only a marginal contribution of within-V1 synaptic inputs. However, our results suggest the possibility that the previously observed link between synaptic inhibition and LFPs may arise from strong co-fluctuations between PV-generated LFPs and the total LFP, as shown by their high R^2^ values (Figure 3F, L; Figure 4E, J; Figure 5E, J). This covariance likely results from the robust drive of PV neurons by external inputs, potentially leading to an overestimation of PV contributions when analyzed with spike-triggered LFP techniques (Figure 3B, H). Our findings therefore indicate that this relationship reflects covariance rather than a substantial contribution to LFP amplitude by PV neurons, and may have led to an inflated significance attributed to PV inputs in previous studies. Notably, these results pertain to mouse V1 during sensory input: in cases with substantially weaker external inputs, PV neurons may play a more pronounced role than observed here.

Our study also highlighted the role of synaptic activity from non-PV V1 populations. Internal LFP (generated exclusively by synaptic activity originating within V1) were predominantly driven by PV interneurons across cortical layers, with excitatory inputs playing only a marginal role. This finding is consistent with earlier reports suggesting that inhibitory perisomatic inputs to pyramidal cells drive LFP and EEG signals, while excitatory inputs contribute more diffusely, resulting in weaker LFP generation ^53,63^. Interestingly, Sst and Htr3a (or equivalently VIP) interneurons contributed negligibly to LFPs, likely due to their lower firing rates and less spatially clustered synaptic activity (Figure S4).

While the contribution of local V1 populations to the overall LFP amplitude was minimal, our results also indicate that their role in affecting the LFP should not be overlooked. Local neuronal processing impacts network firing rates and membrane potential fluctuations, both of which were found to be critical for accurate LFP estimation (Figure 2). This aligns with previous findings that highlighted substantial correlations between LFPs and neuronal membrane potentials ^77,81,82^.

### Limitations

As previously discussed, incorporating non-stationary network dynamics into the kernels introduced a necessary complication to the LFP estimation process, making the kernel-based method parametric. While this extension represents a significant advancement by enabling the kernel method to be applied in more complex, dynamic regimes, it also introduces specific limitations. One notable challenge is the need to pre-define discrete levels for firing rates and the corresponding changes in passive membrane conductance. This requirement adds a layer of complexity, as the number and range of discrete levels must be chosen in advance, which may limit flexibility.

More broadly, this result highlights that accurate LFP estimation cannot be indifferent to the underlying neuronal dynamics of the network. Our results demonstrated that even identical input to the same neuronal populations can yield different LFP patterns, depending on the ongoing neuronal activity.

While this phenomenon may not be entirely surprising, it represents a novel demonstration of the importance of considering the dynamic state of the network. Accounting for these fluctuations is crucial for achieving accurate LFP predictions.

It should be noted that the need to account for fluctuations in firing rates and membrane potentials is not a limitation unique to the kernel-based approach. Persistent pre-synaptic activity alters synaptic driving forces and post-synaptic membrane leak conductances, directly influencing LFP generation. Accordingly, we showed that failing to take into consideration the underlying neuronal dynamics decreased LFP accuracy (Figure 2A vs Figure 2D, and Figure 2B vs Figure 2E). Therefore, the need to account for neuronal dynamics fluctuations over time is a challenge inherent to any framework aiming to accurately predict LFPs in highly non-stationary scenarios. Importantly, such non-stationarities, as observed in the V1 model, reflect the natural variability of biological sensory cortices rather than model flaws. This variability arises from numerous factors, including external input rates ^83,84^, sensory stimulus properties ^85–88^, brain activity patterns ^80^, locomotion ^89^, and transitions between up-down states ^90,91^. We have also shown that the kernel method failed to produce accurate LFP estimations in the presence of substantial radial asymmetries. Specifically, approximately half of the L5 excitatory cells in the model exhibited asymmetrical firing patterns orthogonal to the main axis along which neuronal morphologies are aligned. This, in turn, led to unbalanced LFP fluctuations along the cortical depth (see Figure 6B,E), which the kernel method was not designed to reproduce. However, this increased firing rate along the edge of the L5 population can probably be regarded as an unintended feature of the network, one that might not align with experimentally validated biological scenarios. Consequently, the kernel method’s inability to capture this effect does not represent a significant limitation for most applications.

Finally, we employed the kernel method to disentangle the relative contributions to the V1 LFPs, and we identified the prominence of external inputs. It is important to emphasize that experimental validation is still required to substantiate this claim. Nonetheless, two key considerations suggest the robustness of our findings. First, the V1 computational model used in this study was rigorously validated by incorporating extensive parameter curation and substantial experimental data in its design ^59,66,69^. The so-derived LFP kernels follow deterministically from these parameters. Second, the pronounced effect size of the relative contribution of external synaptic inputs, compared to locally generated inputs, makes it highly plausible that our predictions reflect the biological underlying mechanisms.

### Future developments

The degree to which LFP kernels will be stereotypical or highly variable between different brain regions and species is not currently known, but it is in principle straightforward to investigate. Here we have only focused on mouse V1, where a substantial amount of experimental data is available ^59,66,68,69,92^, but it could be highly valuable to estimate similar kernels also for other often-studied systems in neuroscience. This would allow more researchers to apply the kernel method for calculating LFP signals directly from high-level simulations of neural activity, and it would help us better understand what LFP signals in different brain areas reflect.

One promising direction for future development involves extending the kernel method to account for the temporal evolution of network parameters such as synaptic strengths, particularly those modified by mechanisms like short-term ^93^, spike-timing-dependent ^94^, as well as structural ^95,96^ plasticity. Incorporating these dynamic processes would further refine the method’s predictive capabilities, making it applicable to a wider range of network states and contexts.

Another potential avenue lies in leveraging the kernel method’s formulation, which decomposes LFPs into contributions from different neuronal populations. This approach could be explored to assist in estimating firing rates. In our study, LFPs were indeed derived by starting with known neuronal firing rates and applying spatiotemporal kernels. However, in real-world applications, the typical workflow is reversed: researchers first record extracellular potentials (spikes and/or LFPs) and then aim to infer the underlying neuronal activity that generated these signals. Currently firing rate estimation primarily relies on analyzing the high-frequency components of extracellular potentials (typically above a few hundred Hz), referred to as multi-unit activity (MUA), which reflects the spiking activity of neurons near the electrode. Additionally, spike-sorting algorithms can be then applied to this MUA to disentangle the contributions of (putative) excitatory neurons and fast-spiking interneurons. However, these approaches focus on the microscopic scale (providing information about neurons immediately surrounding the electrode) whereas LFPs capture the (synaptic) activity of larger neuronal populations at the mesoscopic scale.

Current methods of estimating neuronal firing rates often neglect the valuable information embedded in LFPs. Our findings suggest that it may be feasible to use the kernel method to reverse the approach we adopted in this study, utilizing LFP decomposition to assist in estimating the neuronal firing rates from extracellular recordings. Such a methodology could significantly enhance the precision of firing rate estimation by complementing MUA data with the broader mesoscopic information contained in LFPs.

Lastly, another interesting application of the kernel method could be in enhancing the precision of LFP decomposition techniques, such as laminar population analysis ^97^. This tool was recently ^60^ applied to the same network used in this study and was observed to accurately separate the predominant contributions from the different external inputs, as well as the cumulative contribution from recurrent activity within V1.

Incorporating the LFP-kernels into such an analysis could potentially increase its granularity and accuracy, offering a more detailed view of the network’s structure and input dynamics.

## Conclusions

Our study demonstrates the efficacy and versatility of a kernel-based framework for estimating LFPs in large- scale, biophysically detailed neural network models. By integrating dynamic network features such as non- stationary firing rates and fluctuating membrane potentials, we achieved highly accurate and computationally efficient LFP predictions. Importantly, in the mouse V1 model adopted here, we revealed that external inputs, particularly feedback from higher visual areas and thalamic afferents, dominate LFP generation, with minimal contributions from local synaptic activity. This finding not only reinforces the notion of LFPs as signals reflecting network inputs but also identifies their specific origins, offering new insights into cortical processing.

The kernel method’s adaptability, efficiency, and anatomical fidelity pave the way for its application across diverse brain regions and experimental paradigms, and has the potential to advance our understanding of extracellular signals, bridging microscopic and mesoscopic scales of brain activity.

## Methods

### V1 model

In this study, we adopted the V1 mouse model developed by the Allen Institute ^66^, specifically the version presented in (Rimehaug et al., 2023)^59^. For full details, including a thorough description of the data-driven approach justifying the properties and parameter choices, we refer to the original publications ^59,66^.

The V1 model is composed of 230,924 neurons. 51,978 of these are biophysically detailed multicompartment neurons and they form a cylindrical core with diameter of 800 μm and height 860 μm. This inner cylinder is surrounded by an annulus with thickness 445 μm of 178,946 leaky-integrate-and-fire (LIF) point neurons whose purpose is to avoid boundary artifacts ^69^.

From layers 2/3 to 6, there is one excitatory class and three inhibitory classes (PV, Sst, Htr3a) unique to each layer. In layer 1, there is only one Htr3a inhibitory class and no excitatory neurons.

The network received input from three different external sources: (i) experimentally recorded afferent activity from the lateral geniculate nucleus (LGN); (ii) experimentally recorded feedback (FB) from the lateromedial (LM) area of the visual cortex; and (iii) Poisson background spiking activity representing the (stimulus-independent) continuous influence of the rest of the brain on V1.

The LGN module is composed of 17,400 units that are connected to the excitatory classes and the PV interneurons in all layers and the Htr3a class in L1. The FB from LM comprises a single node that provides input to the excitatory, PV, and Sst cells in L2/3 and L5 and represents the feedback from higher visual areas. The background is also a single node, and it is a Poisson source firing at 1 kHz providing input to every neuron in the V1 model.

Recurrent connection probabilities depended on intersomatic distance as well as neurons’ functional preferences, such as direction tuning. Synaptic strengths followed an orientation-dependent like-to-like rule: the connection strength was a function of the difference between the neurons’ preferred orientation and the type of connected neuron pair (i.e., excitatory or inhibitory).

The synaptic location for the MC model between connected neurons depended on the neuronal classes. Excitatory-to-excitatory synapses were placed on the basal and apical dendrites, while avoiding the soma. They were placed anywhere along the dendrites in layers 2/3 and 4, while in layers 5 and 6 they had to be within 200 μm and 150 μm from the soma, respectively. Excitatory to inhibitory synapses were placed on the somas and the basal dendrites.

Inhibitory-to-excitatory and inhibitory-to-inhibitory connections followed instead same targeting rules, which depended on the presynaptic cell type. PV synapses were placed on the soma and dendrites within 50 μm from the soma. Synapses from Sst neurons were placed on the dendrites, 50 μm or further from the soma. Finally, synapses from Htr3a neurons were placed on the dendrites, from 50 μm to 300 μm from the soma.

Thalamic afferents contacted excitatory cell dendrites up to 150 μm away from the soma, excluding the soma itself. LGN synapses onto PV interneurons were placed on the soma and anywhere on the basal dendrites.

FB from LM synapses contacted PV and Sst somas and basal dendrites. FB synapses on L2/3 excitatory cells were placed on the apical dendrites, while on L5 excitatory cells they were placed on both the basal dendrites and apical dendrites more than 300 μm from the soma.

Finally, BKG synapses contacted excitatory cells on basal and apical dendrites closer than 150 μm from the soma, and the soma and basal dendrites of inhibitory cells.

### External stimuli

We adopted two different visual stimuli in this study, as in (Billeh et al., 2020)^66^. The first stimulus involved 3 seconds of drifting gratings: each trial consisted of 500 ms of a gray screen followed by 500 ms of sinusoidal drifting gratings. The gratings were at 80% contrast, with a spatial frequency of 0.04 cycles per degree and a 2 Hz temporal frequency.

The second stimulus involved flashes of light, with each trial consisting of 250 ms of a gray screen, followed by 250 ms of a white screen, and then another 250 ms of a gray screen. The contrast was at 80%. Each trial was presented 10 times.

Notably, the inputs from the LGN in the flash simulations and the FB from LM injected into the V1 model for both stimuli were constructed from spike trains recorded experimentally in these structures during the presentation of the aforementioned stimuli (see ^59,66^). The LGN input in the drifting grating simulation was the same input that was used in the simulations presented in (Billeh et al., 2020)^66^ and was constructed using the FilterNet module that generates LGN spike trains to be used as input for different visual stimuli. The spiking data are publicly available in the Neuropixels dataset https://portal.brain-map.org/circuits-behavior/visual-coding-neuropixels.

### Simulation of extracellular potentials in multicompartmental V1 model

To estimate LFPs in the multicompartmental V1 model we used the model version presented in (Rimehaug et al., 2023)^59^. The code files necessary to run this model can be found in the original publication and are publicly available in Dryad: https://doi.org/10.5061/dryad.k3j9kd5b8.

The model was built, and the simulations carried out with the Brain Modeling Tool Kit (bmtk) software ^98^. Detailed instructions on the network data structures and the simulations codes of the model are provided in (Rimehaug et al., 2023)^59^ and (Billeh et al., 2020)^66^.

Bmtk provides extracellular potential estimates using the so-called line-source approximation as the forward modeling method. In this approach, the extracellular potential generated by a neuronal compartment is approximated to originate from a continuous distribution of the transmembrane currents along a line passing through the central axis of the compartment, within an infinite, homogeneous, isotropic, and linear volume conductor.

Measurement sites in the V1 model were treated as an array of infinitesimally small point electrodes arranged in a straight line, aligned with the main axis of MC cylinder core, and separated by 50 μm.

Local field potentials were finally obtained by low-pass filtering the simulated extracellular potentials with a 4th order low-pass butterworth filter with a cutoff frequency of 150 Hz. We considered only frequencies above 0 Hz by subtracting the mean value in each channel.

Crucially, we found that, irrespective of stimulus, the amplitude of LFPs generated by L2/3 excitatory neurons were significantly larger than those generated by interneurons across the whole cortical depth (Figure 1F). In fact, the contribution to the overall LFPs from synaptic inputs onto every interneuron type (and their associated return currents) was minimal, consistent with what had been previously shown for stellate cells with symmetrically placed synapses ^6,67^ (i.e., a so-called close-field arrangement ^70^). Similarly, we observed that the LFPs generated by excitatory neurons in the deeper cortical layers also dominate those generated by interneurons (Figure S1), further emphasizing the minimal contribution of interneurons to the overall LFPs throughout the cortical depth. Accordingly, in our work, we focused our attention on the LFPs generated by synaptic inputs onto pyramidal cells.

### Estimating spatiotemporal kernels

In this study, we adopted the framework developed by (Hagen et al., 2022) ^55^ to estimate spatiotemporal kernels, which were then used to estimate LFPs in the V1 model outlined above. For full details on the framework, we refer to the original publication.

The kernels represent the averaged spatiotemporal spike-to-signal impulse response functions between a pre- and a postsynaptic neuronal family pair. Crucially, the set of deterministic kernels needed for all connection pathways between pre- and postsynaptic populations X and Y are obtained through a single MC neuron simulation per connection, thereby drastically reducing computational requirements.

Briefly the set of deterministic kernels is obtained through the following process.

1. The postsynaptic population Y is represented by a single typical biophysically detailed multicompartment neuron model. Effectively, the entire postsynaptic neuron population is collapsed into a single neuron with linearized membranes receiving all inputs from every presynaptic population.
2. The dynamics of membrane potentials and synaptic currents across the compartments of the target biophysically detailed neuron are represented using equivalent linear approximations around average membrane potential of the target population, and average firing of presynaptic populations.
3. The average synaptic current density for each connection over the whole postsynaptic target neuron is computed using the information contained in the depth-dependent synaptic density and somatic distribution of target cells along the depth-axis. This step assumes radial homogeneity in synaptic placement and somatic distribution.
4. The computed average synaptic density for all connections between populations X and Y is applied to the target MC neuron representing the whole Y family. This enables the computation of the full set of transmembrane currents in each postsynaptic neuron compartment.
5. The resulting full set of transmembrane currents is filtered to account for synaptic delays between neuronal populations and then combined with forward model matrices to obtain the extracellular potentials at the simulated electrode contact points. This yields the final set of spatiotemporal kernels.

Note that the forward model is modified compared to the classical hybrid scheme to incorporate the distribution of somata in space. Here again, it is assumed that each population is radially symmetric around the vertical z-axis, homogeneous within some radius R, and inhomogeneous along the z-axis.

The kernel estimation process outlined above was originally developed under the assumption (among others) of no distance-dependency for connections in terms of connection probabilities and synaptic conductances. However, the V1 model applied in this study is characterized by both space-dependent and cell-dependent connection probability and synaptic strength, against which the kernel method presented by (Hagen et al., 2022) ^55^ was not validated.

The extracellular potentials were ultimately obtained by convolving the spatiotemporal kernels with the population firing rates. Specifically, for each connection pathway between a presynaptic population X and a target population Y, the extracellular potentials generated by the synaptic activity of X onto Y were obtained by convolving the firing rate of X with the spatiotemporal kernels between X and Y. The total extracellular potentials generated at population X were given by the sum of the extracellular potentials generated by each presynaptic source Y.

For the convolutions, we used the discrete convolution function from numpy, numpy.convolve, with mode=’same’ as described by ^55^. The population firing rates were calculated by counting spikes per time bin of width Δt = 0.1 ms, then dividing by bin width to provide a signal with units of spikes/s.

Local field potentials were obtained by low-pass filtering the simulated extracellular potentials with a 4th order low-pass butterworth filter with a cutoff frequency of 150 Hz. Please note that the kernels-based LFPs goodness-of-fit was relatively robust to variations in the cutoff frequency of the low-pass filter applied during LFP definition, remaining above 0.92 even for cutoffs as high as 500 Hz (Figure S4A).Finally, we considered only frequencies above 0 Hz by subtracting the mean value in each channel.

### Extracting Data from the Allen Institute’s V1 model for kernel application

In order to derive the spatiotemporal kernels to produce LFP estimates, the framework developed by (Hagen et al., 2022) ^55^ necessitates specifying several parameters about the V1 network structure ^59,66^. These parameters were derived from the network files freely available <u>here</u> and <u>here</u>.

Specifically, we derived from the V1 computational model:

- For each connection in the network, we derived the average connection probability, defined as the average number of connections per postsynaptic neuron divided by the numerosity of presynaptic neuronal family.
- Despite the somatic depth distribution being specified in the build .csv files of the computational model, we derived the somatic distributions by fitting uniform probability density functions to the actual cell positions contained in the .h5 network files of the model. This choice was justified by observations that some soma positions did not follow the somatic distribution under which they were built. The reason is that for some cells, if the soma was placed too close to the upper layer boundary, its apical dendrite would protrude out of the pia. Accordingly, the somatic positions were adjusted accounting for the apical dendrite length. Consequently, the actual somatic position distribution was derived from the actual network .h5 files rather than the build .csv files.
- In the original implementation^55^, the kernel code only accounted for gaussian probability density functions of somatic positions. We modified the code to consider uniform somatic positions distributions, as in the case of the V1 model here adopted.
- To compute the kernels, the synaptic distribution along the depth axis onto a representative MC neuron of the target population need to be specified. However, the original V1 network files did not contain any possibility to gather information regarding the synaptic placement onto the neuron morphologies. To extract this information, we built from scratch the structure of the biophysical detailed network using the code patch that can be accessed here: https://github.com/AllenInstitute/bmtk/issues/268.

Once we obtained the synaptic positions in space for each connection pathway, subtracted from the postsynaptic neuron depth, we fit each distribution with a Gaussian mixture model using the GaussianMixture implementation from the sklearn package. The number of components for the fit was chosen to minimize both the Akaike Information Criterion and the Bayesian Information Criterion, with a maximum of 20 components. Goodness-of-fit was computed using the R-squared metric ^99^. Fits with an R-squared value below 0.9 were manually checked and adjusted by increasing the number of components in the Gaussian mixture (this occurred for less than 1% of connections).

- As already anticipated, the kernel method requires a static scalar specifying the connection synaptic weight for each connection pathway. Unfortunately, synaptic strength was not readily available in the network structure files. Consequently, we computed the synaptic strength for each pair of connected neurons in the network. To achieve this, we used the “*DirectionRule_EE*” and the “*DirectionRule_others*” functions developed by the Allen Institute and available <u>here</u>. From the derived strength values, we summarized the distribution of synaptic strengths between neurons of family X contacting those of family Y by computing the median value. We opted for the median value as a measure of central tendency due to the significant asymmetry of synaptic strength distributions (especially between excitatory cells, see Figure S2).
- In the V1 model, each neuron morphology is characterized by its own rotation angle, applied to align the neuron’s main axis with the central axis of the network’s cylindrical core. Unfortunately, the bmtk simulator and the kernel method adopted different x-y-z coordinate systems. Specifically, bmtk assumed the y-axis was aligned with the depth axis, while in the kernel method, the depth axis was aligned with the z-axis. We accounted for this inconsistency between the two frameworks by recomputing the rotation matrices needed to align the neuronal morphology along the depth axis in the kernel method.
- We derived the multapse count (i.e., the number of multiple connections between the same pair of cells) and the synaptic conduction delays from the build .csv files of the V1 network model.

### Single-layer L2/3 V1 model

Before applying the kernel-based LFP estimation to the full-column V1 model, we focused specifically on the upper L2/3 layers. This single-layer model was created by isolating the L2/3 layer within the full-column V1 model. We achieved this by setting the synaptic strengths of connections involving neuronal populations outside L2/3 to zero. This adjustment was made by modifying the csv build files, which specify the synaptic strengths between connected neuronal populations. This allowed us to concentrate solely on the synaptic interactions within the L2/3 layer for a more controlled and detailed analysis.

### Estimating dynamic kernels for LFP estimation

As outlined in the Results section, we proposed a novel method for estimating LFPs based on time-varying kernels. Specifically, we found that the goodness-of-fit of LFP estimates improved when the kernel estimation procedure accounted for temporal fluctuations in two parameters, which were assumed to be constant in the original implementation ^55^: (1) the membrane potential of the neuron representing the target population and (2) the change in passive conductance of the target neuron induced by persistent presynaptic activity.

The first parameter, fluctuations in the membrane potential, was easily handled by applying a linear adjustment, as described in the Results section. However, the second parameter (i.e, changes in passive leak conductance) was more challenging to address analytically and required a different approach.

The concept of incorporating changes in passive leak conductance due to presynaptic activation was already introduced in the original LFP-kernels method ^55^. In their work, the authors calculated changes in the total leak membrane conductance per postsynaptic compartment m of postsynaptic neuron with:

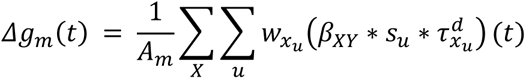

where 𝐴_𝑚_ is the area of compartment m, 𝑠_𝑢_ is the sequence of presynaptic spikes of the unit u belonging to presynaptic population X and contacting the compartment m, 𝜏^𝑑^ the axonal conduction delay, and 𝛽 is the integral of the sum of exponentials describing the rise and the decay of the synaptic conductance dynamics. The asterisk symbol (*) denotes a temporal convolution.

In our work, we used the sum over the compartments of 𝛥*g*_𝑚_ on the neuron representing the postsynaptic population Y, and referred to this as 𝛥*g*_𝑌_(𝑡).

To handle the temporal dynamics of 𝛥*g*_𝑌_(𝑡), we discretized the effective leak conductance into N discrete levels. Specifically, we divided the range of 𝛥*g*_𝑌_(𝑡) into N equal intervals (in our case, N=2) and calculated the average value of 𝛥*g*_𝑌_(𝑡) within each interval.

These N average values represented the discrete levels of 𝛥*g*_𝑌_(𝑡), which we denoted as 𝛥*g*^𝛼^, where α ranges from 1 to N. We then estimated a different set of LFP kernels for each 𝛥*g*^𝛼^. Finally, at each time point, we computed the LFP by adopting the set of kernels corresponding to the discrete value 𝛥*g*^𝛼^ closest to the actual value of 𝛥*g*_𝑌_(𝑡) at that time.

### Developing L2/3 point leaky integrate-and-fire neuron model

As noted in the main text, (Rimehaug et al., 2023)^59^ proposed an updated version of the MC model originally presented by (Billeh et al., 2020)^66^. This new version included additional feedback inputs from the lateromedial visual areas. As a result, however, the synaptic weights of recurrent connections and background inputs were adjusted to ensure consistency in the simulated dynamics between the two model versions. The detailed set of weight adjustments can be found in the original publication^59^.

Unlike (Billeh et al., 2020)^66^, (Rimehaug et al., 2023)^59^ did not develop a point neuron counterpart for the newly proposed version of the MC network model. To address this, we started with the point generalized leaky integrate-and-fire neuron model developed by (Billeh et al., 2020)^66^ and applied the same set of synaptic weight adjustments that (Rimehaug et al., 2023)^59^ applied to the MC model. Please note that our focus was solely on the L2/3 neurons to create a point neuron model equivalent to the single-layer L2/3 MC model.

However, simply applying the same synaptic weight scaling as in (Rimehaug et al., 2023)^59^ was insufficient to ensure consistency between the average firing rates of the L2/3 point network and the L2/3 MC model. To achieve this consistency, the synaptic weights for recurrent connections and external stimuli were multiplied by factors in the range [0.7, 1.3]. The combination of synaptic weights that minimized the difference in average firing rates between the two models, in response to both drifting gratings and flashes of light, was retained (Figure S6) and used for estimating the LFPs with the kernel method and the linear combination of synaptic currents (see ”Layer 2/3 LFP can be computed from a linear combination of synaptic currents simulated with point neuron simulations” in the main text).

### Goodness-of-fit measures

To quantify the temporal agreement between the LFPs obtained through MC model simulations and those obtained via the kernel method we computed the squared of the coefficient of determination R at zero-time lag. The R^2^ was computed as the ratio between the covariance between the two LFP estimates and the product of the variance of each LFP estimate.

In order to quantify instead the relative differences in amplitude between two signals, we computed the ratio the variance between the two signals (see for example Figure 3C-E).

## Resource availability

### Lead contact

The lead contact for this work is Torbjørn V. Ness at.

### Materials availability

This study did not generate new unique reagents.

### Data and code availability

In this work we used two open source freely accessible tools.

The V1 model was originally developed by (Billeh et al., 2020)^66^. In our work we adopted the novel version presented in (Rimehaug et al., 2023)^59^. The files necessary to run simulations of the V1 model used here are publicly available in Dryad: https://doi.org/10.5061/dryad.k3j9kd5b8.

The kernel approach to estimate LFP was originally developed presented by (Hagen et al., 2022)^55^. The codes developing this version of the kernel method are publicly available here: https://github.com/LFPy/LFPykernels.

The codes for the kernel version presented in this code can be found instead here: https://github.com/nicolomeneghetti/LFPkernel_AllenInst_V1model.

## Acknowledgments

The authors wish to thank Anton Arkhipov and Kael Dai for their insightful contributions and unwavering support. Their scientific and technical assistance has been invaluable in the success of this research.

This work received funding from the European Union Horizon 2020 Research and Innovation Programme under Grant Agreement No. 945539 [Human Brain Project (HBP) SGA3], and No. 101147319 [EBRAINS 2.0]. A.M. was supported by the Italian Ministry of Research, under the complementary actions to the NRRP “Fit4MedRob—Fit for Medical Robotics” Grant (# PNC0000007). N.M. was supported by #NEXTGENERATIONEU (NGEU) and funded by the Ministry of University and Research (MUR), National Recovery and Resilience Plan (NRRP), project EBRAINS-Italy (IR0000011) - European Brain ReseArch INfrastructureS-Italy (DN. 101 16.06.2022). The work was funded by “Fondo di beneficenza ed opere di carattere sociale e culturale” granted by “Banca Intesa San Paolo” in the cotext of the project ONDA (“Origini neurali delle disfunzioni dell’arto superiore su modello murino di parkinson”), CUP B53C22007580007.

## Author contributions

Conceptualization: N.M., A.M., G.E., T.V.N.; Methodology: N.M:, T.V.N.; Software: N.M., A.R.; Formal analysis: N.M., Writing – Original Draft, N.M and T.V.N.; Writing – Review & Editing, N.M., A.R., G.E., A.M., T.V.N.; Funding Acquisition, G.E., A.M.; Supervision, G.E., A.M., T.V.N.

## Supplemental information

Document S1. Supplemental text: “LFP proxies based on synaptic currents from point-neuron simulations”. Figures S1–S10.

## Supplemental text

### LFP proxies based on synaptic currents from point-neuron simulations

To compare the kernel method with another well-established simplified LFP prediction approach ^1^, we investigated the feasibility of computing LFPs using standard output from leaky-integrate-and-fire (LIF) network simulations. We designed a point-neuron network equivalent to the multicompartmental L2/3 network discussed previously (see Methods). This point-neuron network, composed of generalized LIF neurons ^2^, shared an identical connectivity graph with the multicompartmental network. To match neuronal dynamics, we adjusted synaptic weights in the point-neuron network to replicate the firing rates observed in the multicompartmental model in response to both stimuli (Figure S6).

To obtain the ground-truth LFPs, we used the previously described LFP kernels (Figure 3), calculating the LFP as the convolution between those kernels and the neuronal firing rates obtained from the point-neuron simulations. This approach highlights that the kernel method can estimate LFPs based on neuronal activity generated from simplified network models, such as point-neurons networks.

Following ^1^, we used non-negative least squares to determine the optimal linear combination of synaptic currents from point-neuron simulations to match the ground-truth LFPs. Since excitatory cells were the primary contributors to LFPs (Figure 1F and Figure S1), we focused on the synaptic currents of excitatory cells.

Our results showed that internal LFPs were accurately approximated by this linear combination of synaptic currents, both for drifting gratings (Figure S7A, median R^2^=0.90) and light flashes (Figure S7B, median R^2^=0.94). To account for changes in passive conductance (𝛥*g*_𝑌_), we optimized separate sets of coefficients for the different conductance levels (N=2). Using static coefficients, instead, reduced the goodness-of-fit for both gratings (median R^2^=0.70) and flashes (median R^2^=0.89).

Consistently with earlier findings (Figure 3), removing synaptic currents from Sst and Htr3a interneurons had minimal impact on LFP accuracy, with a mean R^2^ difference of 0.006 for gratings and 0.004 for flashes. Furthermore, we found the coefficients were stimulus-dependent (Figure S7A compared to Figure S7B), likely reflecting how presynaptic firing rates and membrane potential dynamics influenced LFP generation, and depth-dependent.

Finally, we also found that a linear combination of synaptic currents could approximate LFPs generated by external synaptic inputs, including feedback from LM, background activity, and thalamic afferents (Figure S10). Thus, the overall LFP, comprising both external and internal inputs, was well-approximated by this method (average R^2^=0.92 for gratings, R^2^=0.96 for flashes).

In conclusion, we found that: (1) the kernel method can estimate LFPs from spiking activity in point-neuron simulations; (2) a linear combination of synaptic currents can predict LFPs effectively, even with multiple inhibitory interneuron types; but (3) the synaptic current coefficients depended on cortical depth, stimulus type, and network state, indicating a lack of generalizability, which the kernel method addresses more robustly.

---

**Figure S1.**
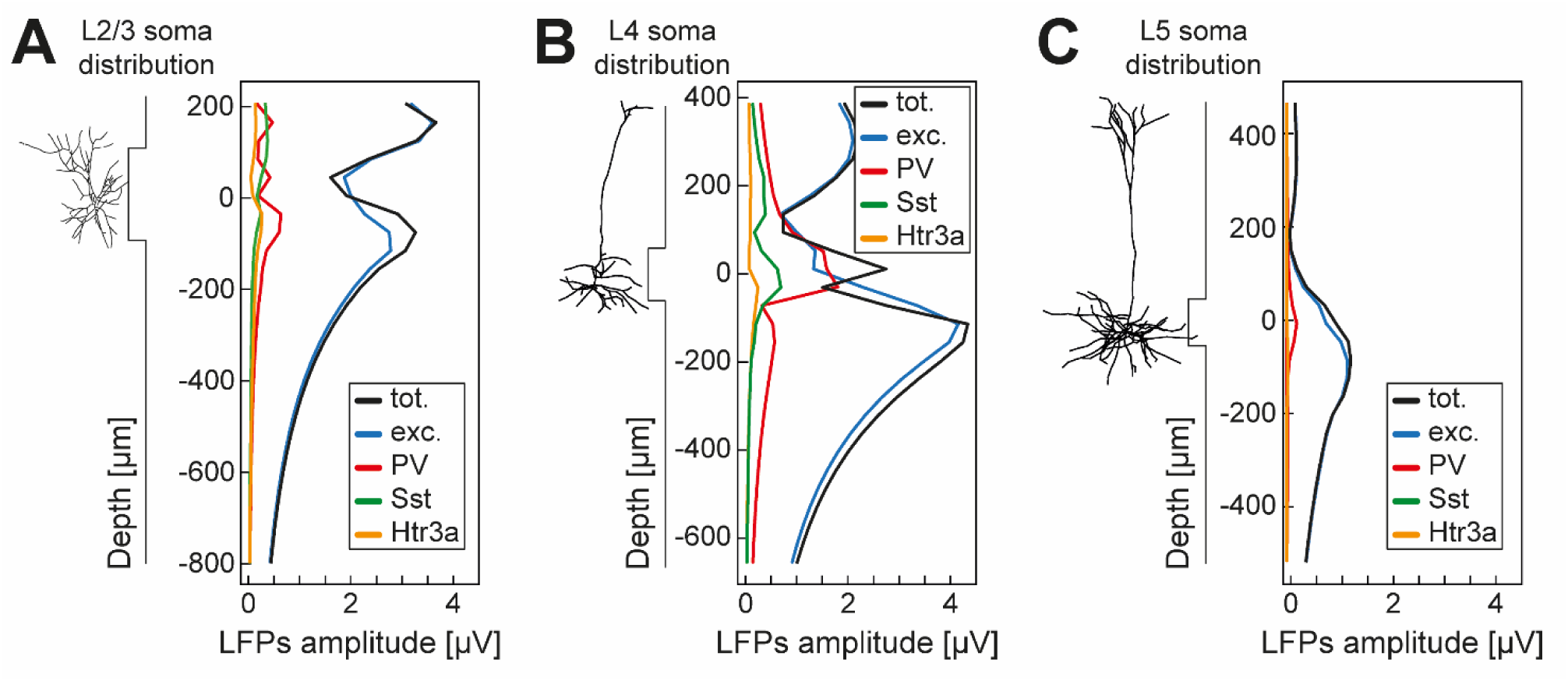
Excitatory cells-generated LFPs predominate over those from inhibitory interneurons. A. Amplitude of LFPs generated at L2/3 neuronal populations in the morphological 3D network across cortical depth. The amplitude was measured as the standard deviation of the LFPs signal over the entire stimulated time course. The amplitudes were averaged between the ones in response to contrast gratings and the ones in response flashes of light. LFPs amplitude was decomposed into the amplitude generated at all neurons (black, ‘tot.’ label), excitatory neurons (blue, ‘exc’), parvalbumin (red, ‘PV’), somatostatin (green, ‘Sst’), and Htr3a interneurons (orange, ‘Htr3a’). The depth axis was centered around the average somatic position of L2/3 cells. B. Same as A, for L4 neuronal populations. C. Same as A, for L5 neuronal populations.

**Figure S2.**
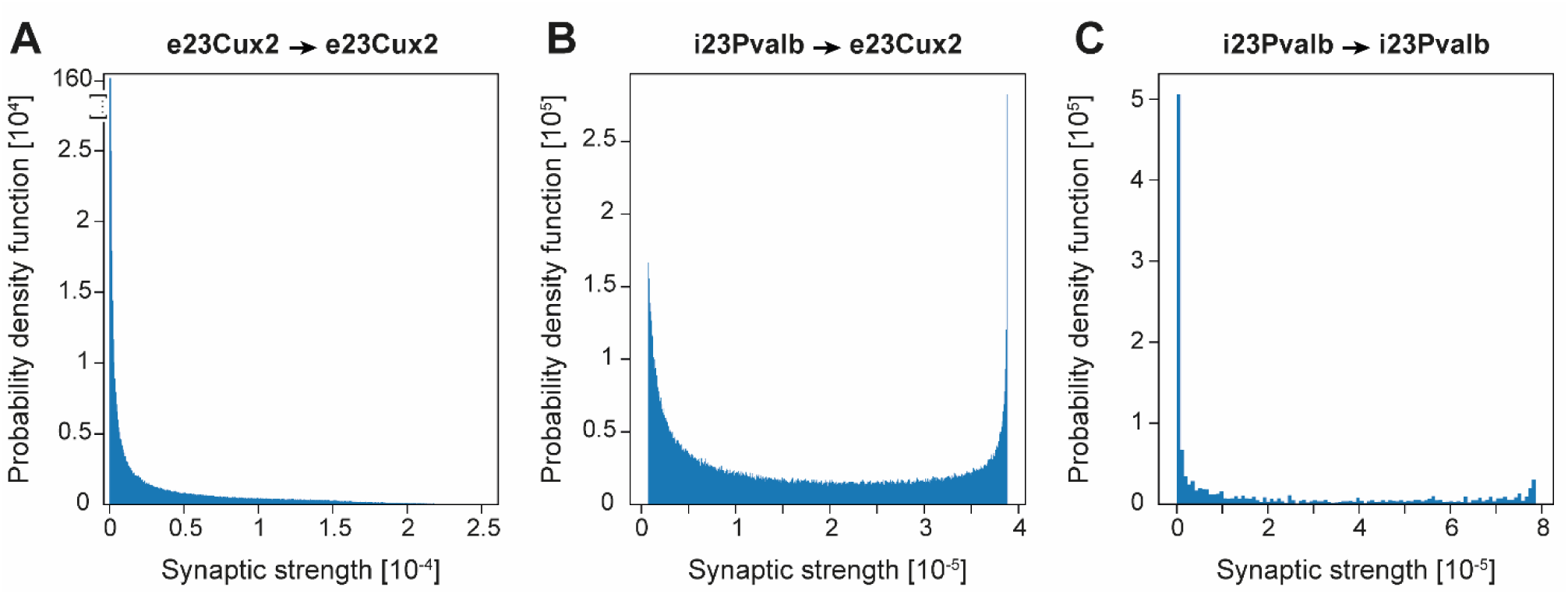
Distribution of recurrent synaptic strengths in V1 model. A. Distribution of synaptic strengths between connected pairs of L2/3 excitatory cells. B. Distribution of synaptic strengths between L2/3 parvalbumin cells and L2/3 excitatory cells. C. Distribution of synaptic strengths between connected pairs of L2/3 parvalbumin cells.

**Figure S3.**
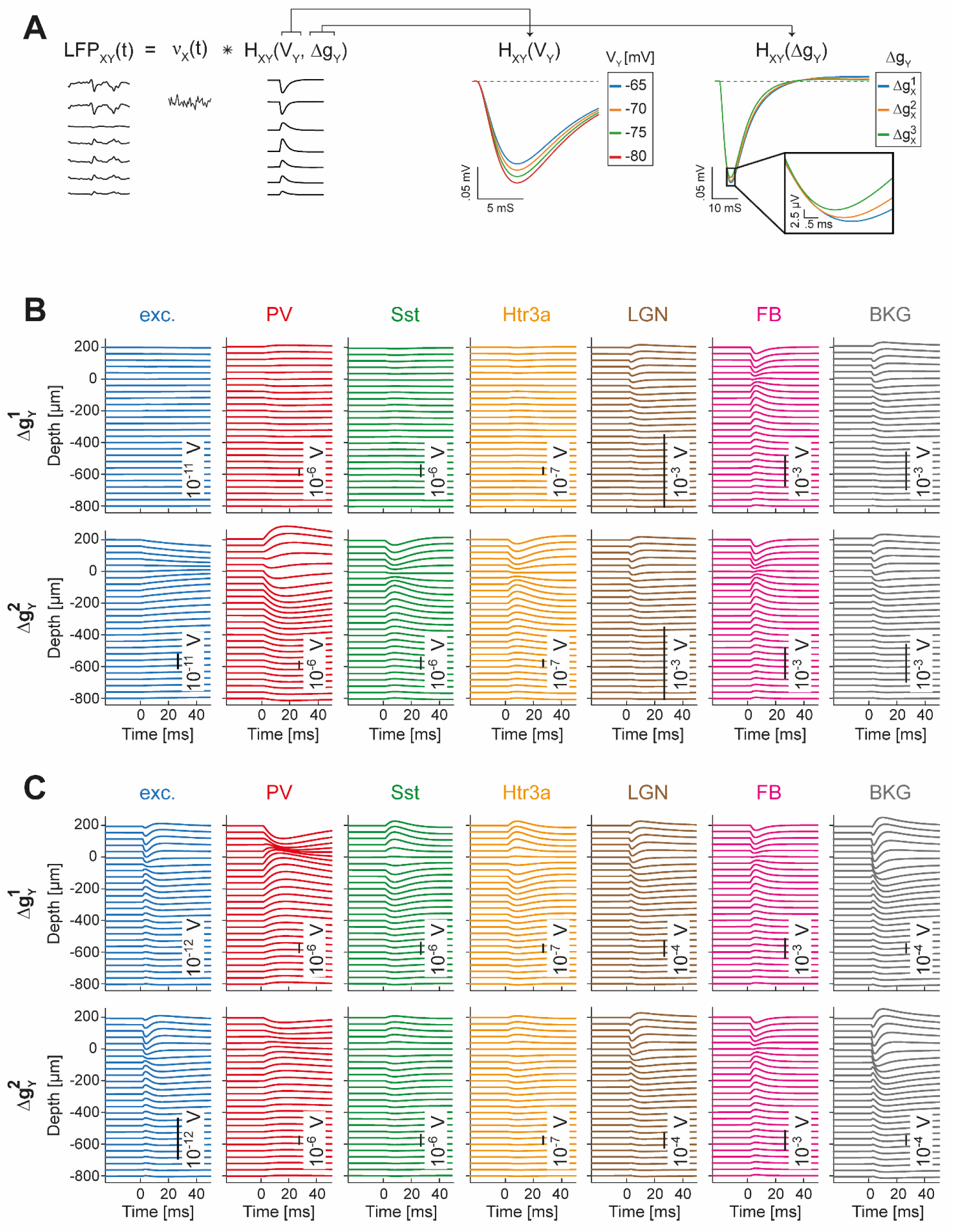
LFP-kernels dynamics for discrete Δg_Y_ levels in response to contrast gratings and light flashes in L2/3 model. A. The LFP_XY_ generated by presynaptic activity from population X onto population Y can be computed as the convolution of the firing rate of X (*υ*_X_) with a set of spatiotemporal kernels (H_XY_). These kernels depend on several factors, including the membrane potential of the target population (V_Y_) and the change in membrane conductance (Δg_Y_) induced by presynaptic activity. The relationship between the kernels and V_Y_ was found to be linear (middle plot shows an example of kernels for four different V_Y_ values). In contrast, the relationship between the kernels and 𝛥*g_Y_* was found to be non-linear (middle plot shows an example of kernels for three different 𝛥*g_Y_* values). B. Average LFP-kernels represented in Figure 3A for both discrete levels of 𝛥*g_Y_* (shown in the two rows) when the single-layer network model was presented with contrast gratings. The sources are (from left to right): excitatory cells (blue); PV (red), Sst (green), and Htr3a (orange) interneurons; LGN (brown), FB (violet), and BKG (grey). Kernel magnitudes are reported in the inset of each panel. C. Same as B, in response to flashes of lights.

**Figure S4.**
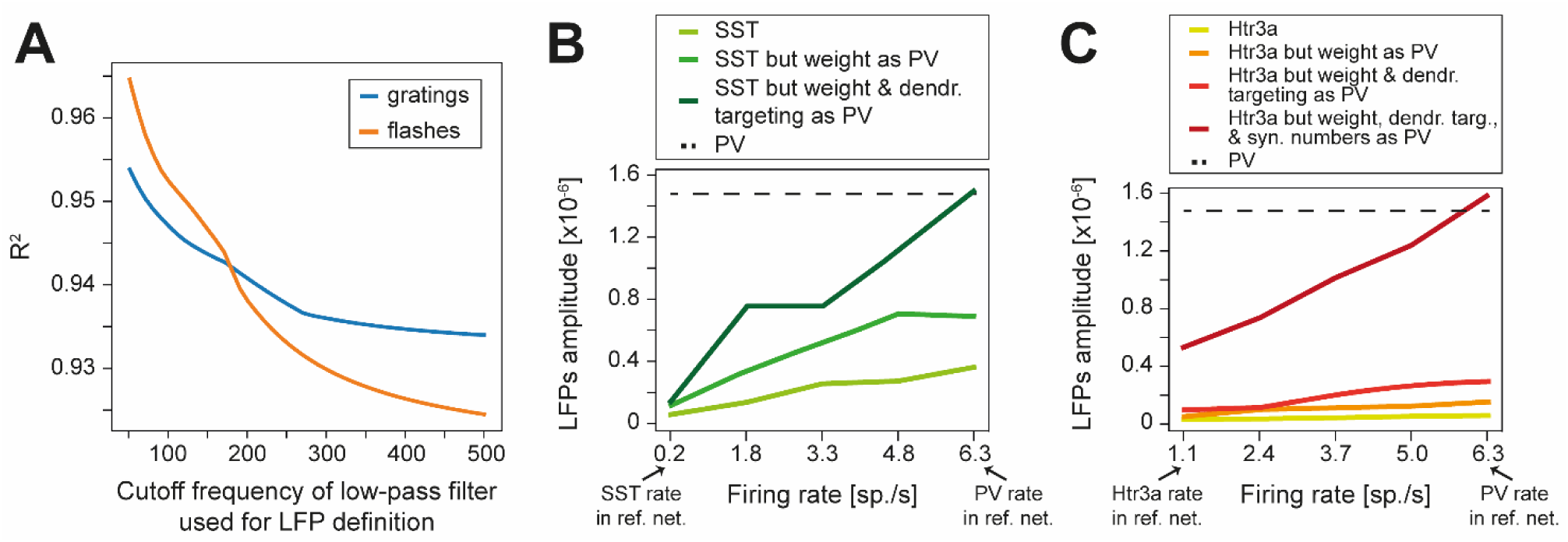
Network properties and neuronal dynamics together contribute to the low LFP amplitude from Sst and Htr3a inhibitory neurons. A. R^2^ between MC and kernel-based LFPs as a function of cutoff frequency used for LFP definition (see Methods) in response to drifting gratings (blue) and flashes of light (orange) for the single-later L2/3 network model. B. LFP amplitude produced by Sst synaptic activity on pyramidal cells within the single-layer L2/3 model. The LFP amplitude was computed based on the simulated average firing rates of Sst neurons under three different conditions: (i) using the same parameters for Sst neurons as in the reference network described in the main text (light green); (ii) applying the same synaptic weight of PV synapses to Sst synapses contacting excitatory cells (green); (iii) as in (ii) but also using the same dendritic targeting as PV synapses (dark green). The simulated firing rates of Sst neurons ranged from their average firing rate in response to contrast gratings in the reference network (i.e., 0.23 sp./s) to the average firing rate of PV neurons in response to the same stimuli (i.e., 6.34 sp./s). The dashed line indicates the LFP amplitude generated by PV interneurons in the reference network (i.e., for a firing rate of 6.34 spikes/s). The amplitude was measured as the standard deviation of the LFPs signal over the entire stimulated time course. The LFPs were simulated by convolving the spatiotemporal kernels across the different conditions with spike trains simulated with a homogenous Poisson process of average firing indicated by the x-axis. The LFP amplitudes reported in the figure represent the average of five simulations. C. Same as B, but for Htr3a interneurons. However, to match the LFPs amplitude generated by Htr3a neurons to those generated by PV interneurons in the reference network (indicated by the dashes line), we computed the LFPs amplitude under four conditions: (i) using the same parameters for Htr3a neurons as in the reference network described in the main text (yellow); (ii) applying the same synaptic weight of PV synapses to Htr3a synapses contacting excitatory cells (orange); (iii) as in (ii) but also using the same dendritic targeting as PV synapses (red); (iv) as in (iii) but also using the same number of synapses per excitatory cell as PV neurons (dark red). The simulated firing rates of Htr3a neurons ranged from their average firing rate in response to contrast gratings in the reference network (i.e., 1.12 sp./s) to the average firing rate of PV neurons in response to the same stimuli (i.e., 6.34 sp./s).

**Figure S5.**
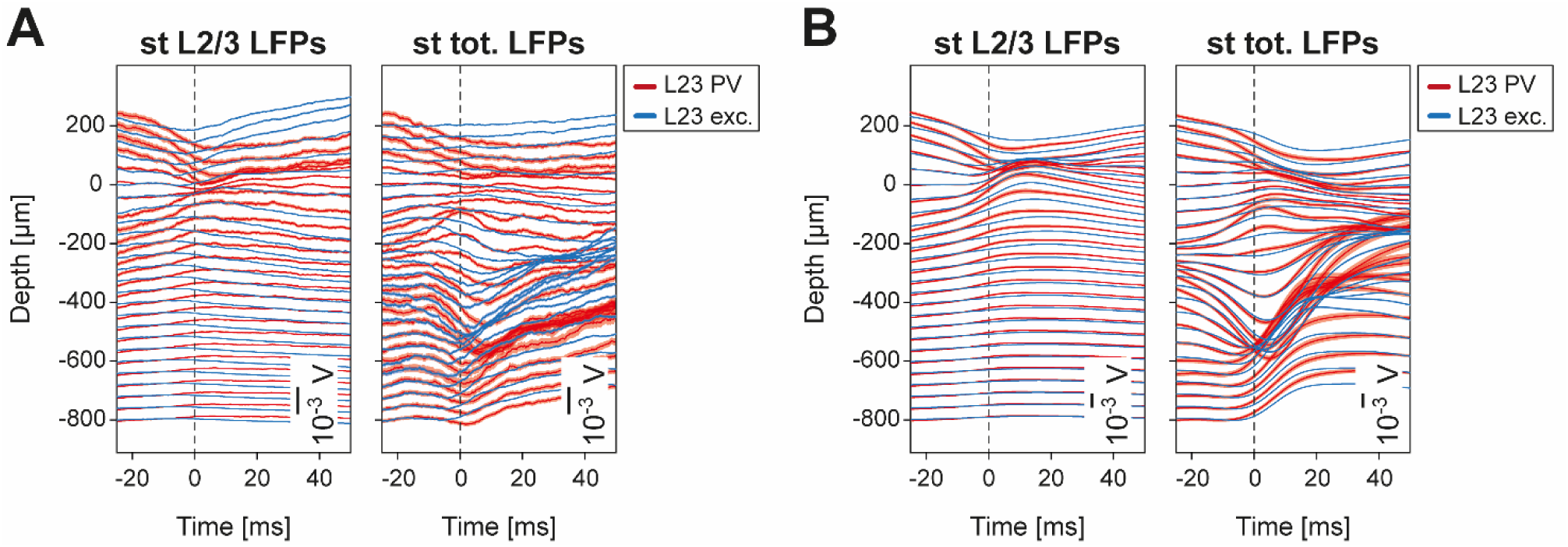
Spike-triggered LFPs by L2/3 PV and excitatory cells. A. (left) Average L2/3 LFPs triggered by spikes of L2/3 PV (red line) or L2/3 excitatory (red line) neurons. Shaded areas indicate the standard error of the mean. L2/3 LFPs indicate the extracellular potentials computed with MC simulations generated by cells within L2/3 in response to drifting gratings in the single-layer V1 model. (right) Average total LFPs triggered by spikes of L2/3 PV (red line) or L2/3 excitatory (red line) neurons. Shaded areas indicate the standard error of the mean. Total LFPs indicate the extracellular potentials computed with MC simulations generated by every cells within the full-column V1 model 3 in response to drifting gratings. B. Same as A, but in response to full-field flashes stimuli.

**Figure S6.**
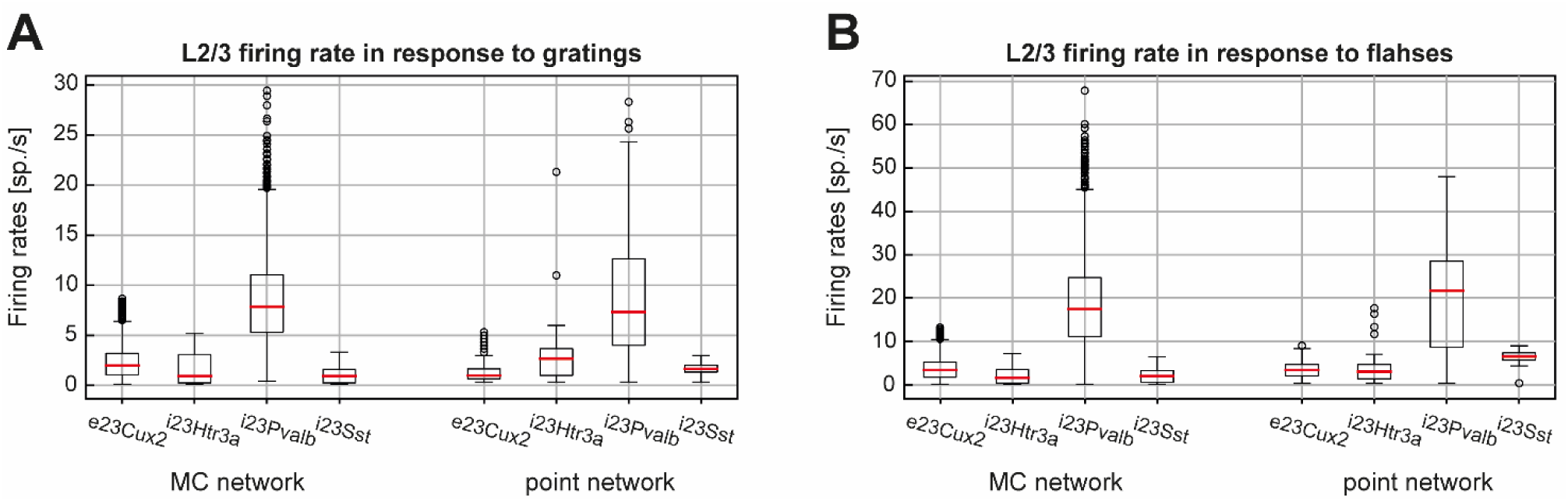
Average firing rate in single-layer L2/3 V1 point-neuron GLIF model. A. Firing rate in response to drifting gratings of L2/3 neuronal populations in MC (4 left boxplots) and point-neuron (4 right boxplots) network simulations. B. Same A, in response to flashes of lights.

**Figure S7.**
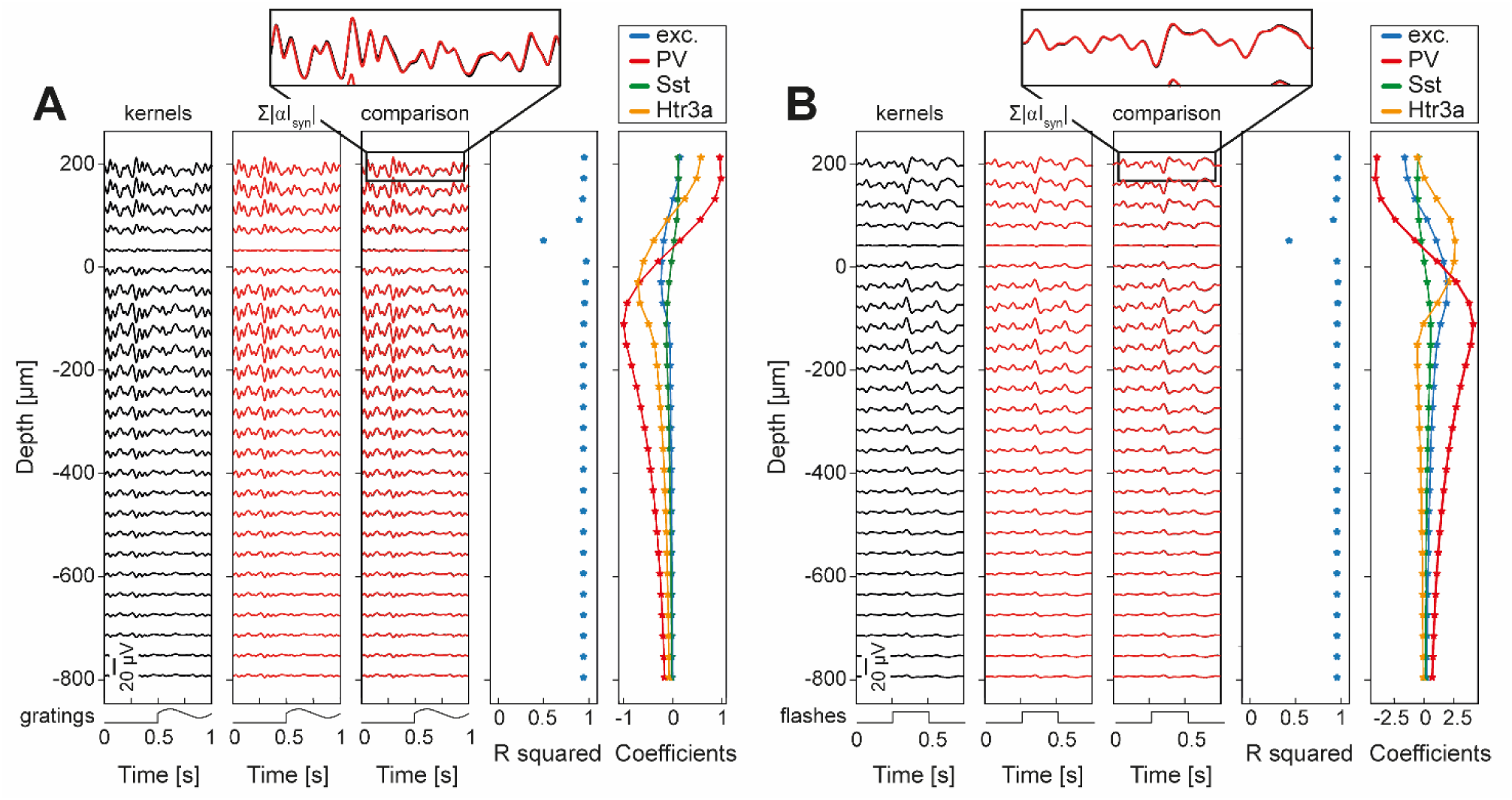
Layer 2/3 LFPs can be computed from the linear combination of synaptic currents simulated with point neuron simulations. A. Depth-resolved trial-averaged L2/3 LFPs in response to visual gratings generated by internal (i.e., within L2/3) neuronal populations (hence excluding external LFPs sources). The temporal trace of the visual stimulus is represented at the bottom of each panel. The LFPs was simulated though a point-neuron simulation as: (i) the convolution between previously described spatiotemporal kernels and neuronal firing rates (black traces, left column); (ii) the linear combination of synaptic currents (red traces, middle column) contacting excitatory cells. The consistency of the two estimation methods was evaluated with R^2^ (fourth column). The linear combination coefficients of synaptic currents for LFPs estimation are reported in the right column: for excitatory cells (blue), PV (red), Sst (green), and for Htr3a (orange) interneurons. The top inset provides a zoomed-in comparison between the LFPs estimated with kernels and with the linear combination of synaptic currents at the highest simulated recording electrode contact. B. Same as A, in response to flashes of light.

**Figure S8.**
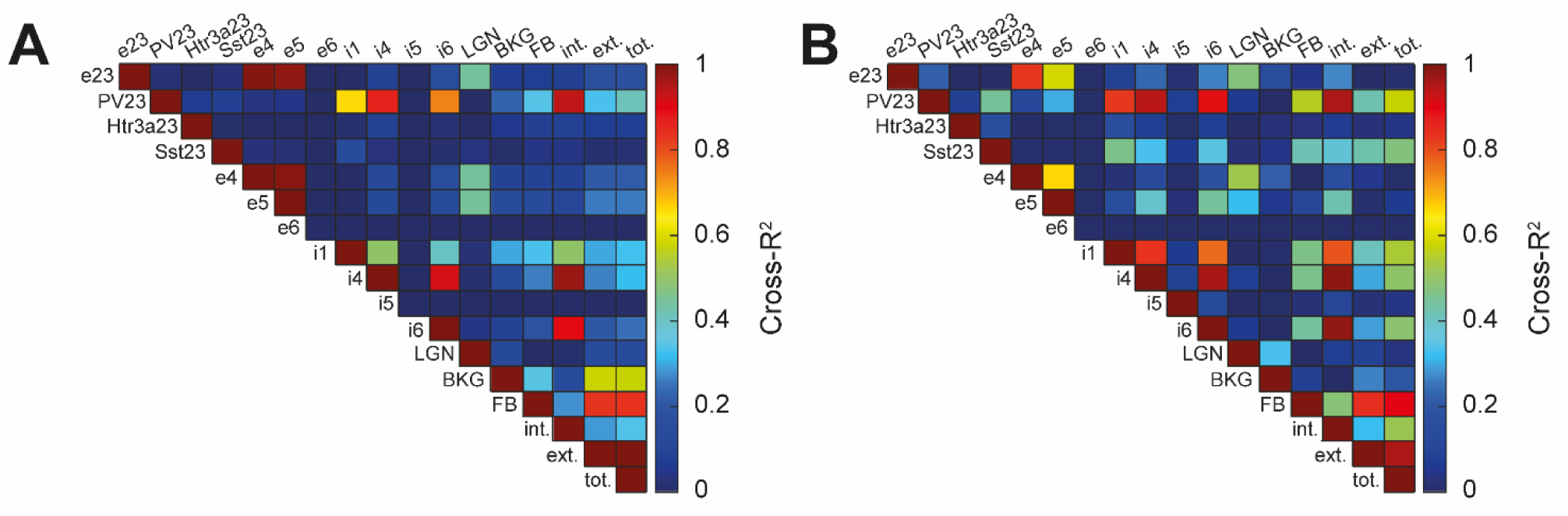
A. R^2^ matrix between the overall L2/3 LFPs and the LFP contributions generated by the firing rates of each neuronal population in the mouse full-column V1 model. . Please note that the labels: ‘intra’ indicate the sum of the LFPs generated by the activity of every internal L2/3 sources; ‘ext’ indicates the sum of the LFPs generated by the activity of every external neuronal family; ‘other I’ indicate the LFPs generated by the activity of inhibitory inputs coming from outside L2/3; ‘other E’ indicates the LFPs generated by the activity of excitatory inputs coming from outside L2/3; ‘tot’ indicates the sum of the LFPs ‘intra’ and ‘ext’. B. Same as A, but in response to flashes of lights.

**Figure S9.**
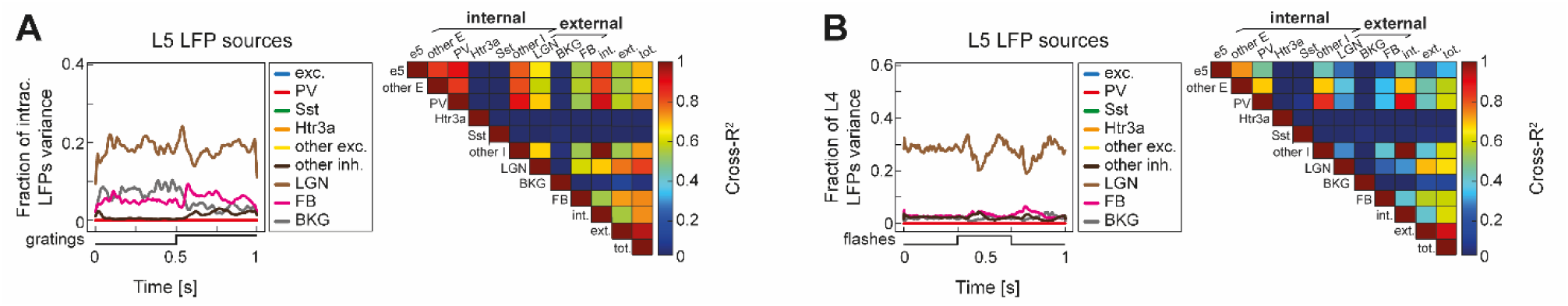
A. (left) Ratio of variance between L5 total LFPs in the mouse V1 full-column model, in response to drifting gratings, and the LFPs generated by the presynaptic activity of: excitatory cells (blue trace); PV interneurons (red trace); Sst interneurons (green trace); Htr3a interneurons (orange trace); inhibitory inputs coming from outside L4 (dark brown trace, labeled ‘other inh.’); excitatory inputs coming from outside L4 (yellow trace, labeled ‘other exc.’); LGN (brown); FB (violet); BKG (grey). (right) R^2^ matrix between the overall L5 LFPs and the LFP contributions generated by the firing rates of each neuronal population in the mouse full-column V1 model. Please note that the labels: ‘int.’ indicate the sum of the LFPs generated by the activity of every V1 populations; ‘ext.’ indicates the sum of the LFPs generated by the synaptic activity of the three external stimuli; ‘other I’ indicate the LFPs generated by the activity of inhibitory inputs coming from outside L5; ‘other E’ indicates the LFPs generated by the activity of excitatory inputs coming from outside L5; ‘tot.’ indicates the sum of the LFPs ‘int.’ and ‘ext.’. B. Same as A, but in response to full-field flashes of lights.

**Figure S10.**
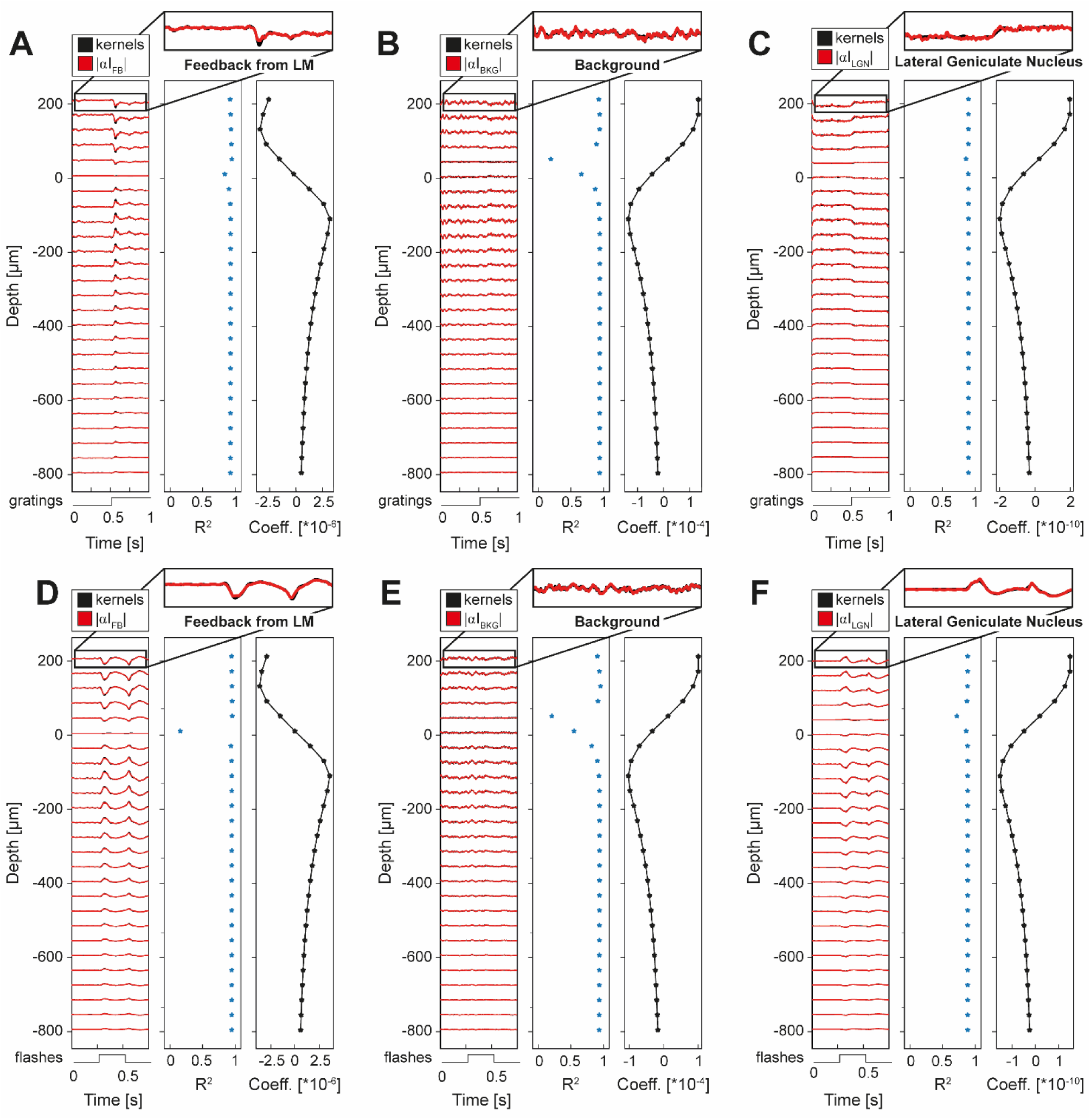
LFPs generated by external inputs in L2/3 point-neuron model can be computed either with kernels or as linear combination of synaptic currents. A. Depth-resolved trial-averaged L2/3 LFPs in response to visual gratings generated by feedback from LM. The temporal trace of the visual stimulus is represented at the bottom of each panel. The LFPs was simulated through a point-neuron simulation as: (i) the convolution between previously described spatiotemporal kernels and neuronal firing rates (black traces); (ii) the linear combination of FB synaptic currents (red traces) contacting excitatory cells. The consistency of the two estimation methods was evaluated with R^2^ (middle column, median R^2^=0.93). The linear combination coefficients across cortical depth for the FB synaptic currents are reported in the right column. The top inset provides a zoomed-in comparison between the LFPs estimated with kernels and with the linear combination of synaptic currents at the highest simulated recording electrode contact. B. Same as A, but for the L2/2 LFPs generated by Poisson background input in response to visual gratings. Median R^2^=0.94. C. Same as A, but for the L2/2 LFPs generated by thalamic afferents in response to visual gratings. Median R^2^=0.94. D. Same as A, but in response to flashes of lights. Median R^2^=0.95. E. Same as B, but in response to flashes of lights. Median R^2^=0.94. F. Same as B, but in response to flashes of lights. Median R^2^=0.91.

## References

1. Einevoll, G.T., Kayser, C., Logothetis, N.K., and Panzeri, S. (2013). Modelling and analysis of local field potentials for studying the function of cortical circuits. Nat Rev Neurosci 14, 770–785. 10.1038/nrn3599.

2. 2. Halnes, G., Ness, T.V., Næss, S., Hagen, E., Pettersen, K.H., and Einevoll, G. (2024). Electric brain signals: foundations and applications of biophysical modeling (Cambridge University Press).

3. Buzsáki, G., Anastassiou, C.A., and Koch, C. (2012). The origin of extracellular fields and currents — EEG, ECoG, LFP and spikes. Nat Rev Neurosci 13, 407–420. 10.1038/nrn3241.

4. Katzner, S., Nauhaus, I., Benucci, A., Bonin, V., Ringach, D.L., and Carandini, M. (2009). Local Origin of Field Potentials in Visual Cortex. Neuron 61, 35–41. 10.1016/j.neuron.2008.11.016.

5. Kajikawa, Y., and Schroeder, C.E. (2011). How Local Is the Local Field Potential? Neuron 72, 847–858. 10.1016/j.neuron.2011.09.029.

6. Lindén, H., Tetzlaff, T., Potjans, T.C., Pettersen, K.H., Grün, S., Diesmann, M., and Einevoll, G.T. (2011). Modeling the Spatial Reach of the LFP. Neuron 72, 859–872. 10.1016/j.neuron.2011.11.006.

7. Łęski, S., Lindén, H., Tetzlaff, T., Pettersen, K.H., and Einevoll, G.T. (2013). Frequency Dependence of Signal Power and Spatial Reach of the Local Field Potential. PLoS Comput Biol 9, e1003137. 10.1371/journal.pcbi.1003137.

8. Dubey, A., and Ray, S. (2016). Spatial spread of local field potential is band-pass in the primary visual cortex. Journal of Neurophysiology 116, 1986–1999. 10.1152/jn.00443.2016.

9. Jun, J.J., Steinmetz, N.A., Siegle, J.H., Denman, D.J., Bauza, M., Barbarits, B., Lee, A.K., Anastassiou, C.A., Andrei, A., Aydın, Ç., et al. (2017). Fully integrated silicon probes for high-density recording of neural activity. Nature 551, 232–236. 10.1038/nature24636.

10. Klein, L., Pothof, F., Raducanu, B.C., Klon-Lipok, J., Shapcott, K.A., Musa, S., Andrei, A., Aarts, A.A., Paul, O., Singer, W., et al. (2020). High-density electrophysiological recordings in macaque using a chronically implanted 128-channel passive silicon probe. J. Neural Eng. 17, 026036. 10.1088/1741-2552/ab8436.

11. Ding, X.-F., Gao, Y., Zhang, H., Zhang, Y., Wang, S.-X., Zhao, Y.-Q., Wang, Y.-Z., and Fan, M. (2020). A novel low-cost electrode for recording the local field potential of freely moving rat’s brain. Translational Neuroscience 11, 96–104. 10.1515/tnsci-2020-0104.

12. Mendoza-Halliday, D., Major, A.J., Lee, N., Lichtenfeld, M.J., Carlson, B., Mitchell, B., Meng, P.D., Xiong, Y., Westerberg, J.A., Jia, X., et al. (2024). A ubiquitous spectrolaminar motif of local field potential power across the primate cortex. Nat Neurosci 27, 547–560. 10.1038/s41593-023-01554-7.

13. 13. Mehring, C., Rickert, J., Vaadia, E., De Oliveira, S.C., Aertsen, A., and Rotter, S. (2003). Inference of hand movements from local field potentials in monkey motor cortex. Nat Neurosci 6, 1253–1254. 10.1038/nn1158.

14. Stavisky, S.D., Kao, J.C., Nuyujukian, P., Ryu, S.I., and Shenoy, K.V. (2015). A high performing brain– machine interface driven by low-frequency local field potentials alone and together with spikes. J. Neural Eng. 12, 036009. 10.1088/1741-2560/12/3/036009.

15. Mitzdorf, U. (1985). Current source-density method and application in cat cerebral cortex: investigation of evoked potentials and EEG phenomena. Physiological Reviews 65, 37–100. 10.1152/physrev.1985.65.1.37.

16. Kandel, A., and Buzsáki, G. (1997). Cellular-synaptic generation of sleep spindles, spike-and-wave discharges, and evoked thalamocortical responses in the neocortex of the rat. J Neurosci 17, 6783–6797. 10.1523/JNEUROSCI.17-17-06783.1997.

17. Schroeder, C. (1998). A spatiotemporal profile of visual system activation revealed by current source density analysis in the awake macaque. Cerebral Cortex 8, 575–592. 10.1093/cercor/8.7.575.

18. Belitski, A., Gretton, A., Magri, C., Murayama, Y., Montemurro, M.A., Logothetis, N.K., and Panzeri, S. (2008). Low-Frequency Local Field Potentials and Spikes in Primary Visual Cortex Convey Independent Visual Information. Journal of Neuroscience 28, 5696–5709. 10.1523/JNEUROSCI.0009-08.2008.

19. Belitski, A., Panzeri, S., Magri, C., Logothetis, N.K., and Kayser, C. (2010). Sensory information in local field potentials and spikes from visual and auditory cortices: time scales and frequency bands. J Comput Neurosci 29, 533–545. 10.1007/s10827-010-0230-y.

20. Mazzoni, A., Brunel, N., Cavallari, S., Logothetis, N.K., and Panzeri, S. (2011). Cortical dynamics during naturalistic sensory stimulations: Experiments and models. Journal of Physiology-Paris 105, 2–15. 10.1016/j.jphysparis.2011.07.014.

21. Barbieri, F., Mazzoni, A., Logothetis, N.K., Panzeri, S., and Brunel, N. (2014). Stimulus Dependence of Local Field Potential Spectra: Experiment versus Theory. Journal of Neuroscience 34, 14589–14605. 10.1523/JNEUROSCI.5365-13.2014.

22. Vinck, M., Batista-Brito, R., Knoblich, U., and Cardin, J.A. (2015). Arousal and Locomotion Make Distinct Contributions to Cortical Activity Patterns and Visual Encoding. Neuron 86, 740–754. 10.1016/j.neuron.2015.03.028.

23. Meneghetti, N., Cerri, C., Tantillo, E., Vannini, E., Caleo, M., and Mazzoni, A. (2021). Narrow and Broad γ Bands Process Complementary Visual Information in Mouse Primary Visual Cortex. eNeuro 8, ENEURO.0106-21.2021. 10.1523/ENEURO.0106-21.2021.

24. Donoghue, J.P., Sanes, J.N., Hatsopoulos, N.G., and Gaál, G. (1998). Neural Discharge and Local Field Potential Oscillations in Primate Motor Cortex During Voluntary Movements. Journal of Neurophysiology 79, 159–173. 10.1152/jn.1998.79.1.159.

25. Scherberger, H., Jarvis, M.R., and Andersen, R.A. (2005). Cortical Local Field Potential Encodes Movement Intentions in the Posterior Parietal Cortex. Neuron 46, 347–354. 10.1016/j.neuron.2005.03.004.

26. Brown, P., and Williams, D. (2005). Basal ganglia local field potential activity: Character and functional significance in the human. Clinical Neurophysiology 116, 2510–2519. 10.1016/j.clinph.2005.05.009.

27. 27. Yun, K., Lebedev, M., and Nicolelis, M.A.L. (2007). Prediction of motor timing using nonlinear analysis of local field potentials. In World Congress on Medical Physics and Biomedical Engineering 2006 IFMBE Proceedings., R. Magjarevic and J. H. Nagel, eds. (Springer Berlin Heidelberg), pp. 1005–1008. 10.1007/978-3-540-36841-0_239.

28. 28. Combrisson, E., Di Rienzo, F., Saive, A.-L., Perrone-Bertolotti, M., Soto, J.L.P., Kahane, P., Lachaux, J.- P., Guillot, A., and Jerbi, K. (2024). Human local field potentials in motor and non-motor brain areas encode upcoming movement direction. Commun Biol 7, 506. 10.1038/s42003-024-06151-3.

29. Asher, E.E., Slovik, M., Mitelman, R., Bergman, H., Havlin, S., and Moshel, S. (2024). Local field potential journey into the Basal Ganglia. Deep Brain Stimulation 5, 20–29. 10.1016/j.jdbs.2024.03.002.

30. Pesaran, B., Pezaris, J.S., Sahani, M., Mitra, P.P., and Andersen, R.A. (2002). Temporal structure in neuronal activity during working memory in macaque parietal cortex. Nat Neurosci 5, 805–811. 10.1038/nn890.

31. Kreiman, G., Hung, C.P., Kraskov, A., Quiroga, R.Q., Poggio, T., and DiCarlo, J.J. (2006). Object Selectivity of Local Field Potentials and Spikes in the Macaque Inferior Temporal Cortex. Neuron 49, 433–445. 10.1016/j.neuron.2005.12.019.

32. Liu, J., and Newsome, W.T. (2006). Local Field Potential in Cortical Area MT: Stimulus Tuning and Behavioral Correlations. Journal of Neuroscience 26, 7779–7790. 10.1523/JNEUROSCI.5052-05.2006.

33. Liebe, S., Hoerzer, G.M., Logothetis, N.K., and Rainer, G. (2012). Theta coupling between V4 and prefrontal cortex predicts visual short-term memory performance. Nat Neurosci 15, 456–462. 10.1038/nn.3038.

34. Doucet, G., Gulli, R.A., Corrigan, B.W., Duong, L.R., and Martinez-Trujillo, J.C. (2020). Modulation of local field potentials and neuronal activity in primate hippocampus during saccades. Hippocampus 30, 192–209. 10.1002/hipo.23140.

35. Prakash, S.S., Mayo, J.P., and Ray, S. (2022). Decoding of attentional state using local field potentials. Current Opinion in Neurobiology 76, 102589. 10.1016/j.conb.2022.102589.

36. Meneghetti, N., Vannini, E., and Mazzoni, A. (2024). Rodents’ visual gamma as a biomarker of pathological neural conditions. The Journal of Physiology 602, 1017–1048. 10.1113/JP283858.

37. Petrucco, L., Pracucci, E., Brondi, M., Ratto, G.M., and Landi, S. (2017). Epileptiform activity in the mouse visual cortex interferes with cortical processing in connected areas. Sci Rep 7, 40054. 10.1038/srep40054.

38. Panarese, A., Vissani, M., Meneghetti, N., Vannini, E., Cracchiolo, M., Micera, S., Caleo, M., Mazzoni, A., and Restani, L. (2022). Disruption of layer-specific visual processing in a model of focal neocortical epilepsy. Cerebral Cortex, bhac335. 10.1093/cercor/bhac335.

39. Gill, B.J.A., Wu, X., Khan, F.A., Sosunov, A.A., Liou, J., Dovas, A., Eissa, T.L., Banu, M.A., Bateman, L.M., McKhann, G.M., et al. (2020). Ex vivo multi-electrode analysis reveals spatiotemporal dynamics of ictal behavior at the infiltrated margin of glioma. Neurobiology of Disease 134, 104676. 10.1016/j.nbd.2019.104676.

40. 40. Tantillo, E., Scalera, M., De Santis, E., Meneghetti, N., Cerri, C., Menicagli, M., Mazzoni, A., Costa, M., Mazzanti, C.M., Vannini, E., et al. (2023). Molecular changes underlying decay of sensory responses and enhanced seizure propensity in peritumoral neurons. Neuro-Oncology, noad035. 10.1093/neuonc/noad035.

41. Krishna, S., Choudhury, A., Keough, M.B., Seo, K., Ni, L., Kakaizada, S., Lee, A., Aabedi, A., Popova, G., Lipkin, B., et al. (2023). Glioblastoma remodelling of human neural circuits decreases survival. Nature 617, 599–607. 10.1038/s41586-023-06036-1.

42. Welle, C.G. (2010). Gamma Oscillations in the Mouse Primary Visual Cortex as an Endophenotype of Schizophrenia.

43. Hamm, J.P., Peterka, D.S., Gogos, J.A., and Yuste, R. (2017). Altered Cortical Ensembles in Mouse Models of Schizophrenia. Neuron 94, 153–167.e8. 10.1016/j.neuron.2017.03.019.

44. Meneghetti, N., Cerri, C., Vannini, E., Tantillo, E., Tottene, A., Pietrobon, D., Caleo, M., and Mazzoni, A. (2022). Synaptic alterations in visual cortex reshape contrast-dependent gamma oscillations and inhibition-excitation ratio in a genetic mouse model of migraine. J Headache Pain 23, 125. 10.1186/s10194-022-01495-9.

45. Adaikkan, C., Wang, J., Abdelaal, K., Middleton, S.J., Bozzelli, P.L., Wickersham, I.R., McHugh, T.J., and Tsai, L.-H. (2022). Alterations in a cross-hemispheric circuit associates with novelty discrimination deficits in mouse models of neurodegeneration. Neuron 110, 3091–3105.e9. 10.1016/j.neuron.2022.07.023.

46. Kühn, A.A., Kupsch, A., Schneider, G., and Brown, P. (2006). Reduction in subthalamic 8–35 Hz oscillatory activity correlates with clinical improvement in Parkinson’s disease. Eur J of Neuroscience 23, 1956–1960. 10.1111/j.1460-9568.2006.04717.x.

47. Telkes, I., Viswanathan, A., Jimenez-Shahed, J., Abosch, A., Ozturk, M., Gupte, A., Jankovic, J., and Ince, N.F. (2018). Local field potentials of subthalamic nucleus contain electrophysiological footprints of motor subtypes of Parkinson’s disease. Proc. Natl. Acad. Sci. U.S.A. 115. 10.1073/pnas.1810589115.

48. Wiest, C., Tinkhauser, G., Pogosyan, A., Bange, M., Muthuraman, M., Groppa, S., Baig, F., Mostofi, A., Pereira, E.A., Tan, H., et al. (2020). Local field potential activity dynamics in response to deep brain stimulation of the subthalamic nucleus in Parkinson’s disease. Neurobiology of Disease 143, 105019. 10.1016/j.nbd.2020.105019.

49. Pettersen, K.H., Hagen, E., and Einevoll, G.T. (2008). Estimation of population firing rates and current source densities from laminar electrode recordings. J Comput Neurosci 24, 291–313. 10.1007/s10827-007-0056-4.

50. Lindén, H., Pettersen, K.H., and Einevoll, G.T. (2010). Intrinsic dendritic filtering gives low-pass power spectra of local field potentials. J Comput Neurosci 29, 423–444. 10.1007/s10827-010-0245-4.

51. Lindén, H., Hagen, E., Łęski, S., Norheim, E.S., Pettersen, K.H., and Einevoll, G.T. (2014). LFPy: a tool for biophysical simulation of extracellular potentials generated by detailed model neurons. Front. Neuroinform. 7. 10.3389/fninf.2013.00041.

52. Reimann, M.W., Anastassiou, C.A., Perin, R., Hill, S.L., Markram, H., and Koch, C. (2013). A Biophysically Detailed Model of Neocortical Local Field Potentials Predicts the Critical Role of Active Membrane Currents. Neuron 79, 375–390. 10.1016/j.neuron.2013.05.023.

53. 53. Hagen, E., Dahmen, D., Stavrinou, M.L., Lindén, H., Tetzlaff, T., van Albada, S.J., Grün, S., Diesmann, M., and Einevoll, G.T. (2016). Hybrid Scheme for Modeling Local Field Potentials from Point-Neuron Networks. Cereb. Cortex 26, 4461–4496. 10.1093/cercor/bhw237.

54. Hagen, E., Næss, S., Ness, T.V., and Einevoll, G.T. (2018). Multimodal Modeling of Neural Network Activity: Computing LFP, ECoG, EEG, and MEG Signals With LFPy 2.0. Front Neuroinform *12*, 92. 10.3389/fninf.2018.00092.

55. Hagen, E., Magnusson, S.H., Ness, T.V., Halnes, G., Babu, P.N., Linssen, C., Morrison, A., and Einevoll, G.T. (2022). Brain signal predictions from multi-scale networks using a linearized framework. PLoS Comput Biol 18, e1010353. 10.1371/journal.pcbi.1010353.

56. Ness, T.V., Remme, M.W.H., and Einevoll, G.T. (2018). h-Type Membrane Current Shapes the Local Field Potential from Populations of Pyramidal Neurons. J. Neurosci. 38, 6011–6024. 10.1523/JNEUROSCI.3278-17.2018.

57. 57. Einevoll, G.T., Destexhe, A., Diesmann, M., Grün, S., Jirsa, V., De Kamps, M., Migliore, M., Ness, T.V., Plesser, H.E., and Schürmann, F. (2019). The Scientific Case for Brain Simulations. Neuron 102, 735– 744. 10.1016/j.neuron.2019.03.027.

58. Haufler, D., Ito, S., Koch, C., and Arkhipov, A. (2023). Simulations of cortical networks using spatially extended conductance-based neuronal models. The Journal of Physiology 601, 3123–3139. 10.1113/JP284030.

59. Rimehaug, A.E., Stasik, A.J., Hagen, E., Billeh, Y.N., Siegle, J.H., Dai, K., Olsen, S.R., Koch, C., Einevoll, G.T., and Arkhipov, A. (2023). Uncovering circuit mechanisms of current sinks and sources with biophysical simulations of primary visual cortex. eLife 12, e87169. 10.7554/eLife.87169.

60. Rimehaug, A.E., Dale, A.M., Arkhipov, A., and Einevoll, G.T. (2024). Uncovering population contributions to the extracellular potential in the mouse visual system using Laminar Population Analysis. Preprint, 10.1101/2024.01.15.575805 10.1101/2024.01.15.575805.

61. Tharayil, J., Blanco Alonso, J., Farcito, S., Lloyd, B., Romani, A., Boci, E., Cassara, A., Schürmann, F., Neufeld, E., Kuster, N., et al. (2024). BlueRecording: A pipeline for the efficient calculation of extracellular recordings in large-scale neural circuit models. Preprint, 10.1101/2024.05.14.591849 10.1101/2024.05.14.591849.

62. Senk, J., Hagen, E., Van Albada, S.J., and Diesmann, M. (2024). Reconciliation of weak pairwise spike– train correlations and highly coherent local field potentials across space. Cerebral Cortex 34, bhae405. 10.1093/cercor/bhae405.

63. Mazzoni, A., Lindén, H., Cuntz, H., Lansner, A., Panzeri, S., and Einevoll, G.T. (2015). Computing the Local Field Potential (LFP) from Integrate-and-Fire Network Models. PLoS Comput Biol 11, e1004584. 10.1371/journal.pcbi.1004584.

64. Telenczuk, B., Telenczuk, M., and Destexhe, A. (2020). A kernel-based method to calculate local field potentials from networks of spiking neurons. J Neurosci Methods 344, 108871. 10.1016/j.jneumeth.2020.108871.

65. Ness, T.V., Tetzlaff, T., Einevoll, G.T., and Dahmen, D. (2024). On the validity of electric brain signal predictions based on population firing rates. Preprint, 10.1101/2024.07.10.602833 10.1101/2024.07.10.602833.

66. Billeh, Y.N., Cai, B., Gratiy, S.L., Dai, K., Iyer, R., Gouwens, N.W., Abbasi-Asl, R., Jia, X., Siegle, J.H., Olsen, S.R., et al. (2020). Systematic Integration of Structural and Functional Data into Multi-scale Models of Mouse Primary Visual Cortex. Neuron, S0896627320300672. 10.1016/j.neuron.2020.01.040.

67. Næss, S., Halnes, G., Hagen, E., Hagler, D.J., Dale, A.M., Einevoll, G.T., and Ness, T.V. (2021). Biophysically detailed forward modeling of the neural origin of EEG and MEG signals. NeuroImage 225, 117467. 10.1016/j.neuroimage.2020.117467.

68. Gouwens, N.W., Sorensen, S.A., Berg, J., Lee, C., Jarsky, T., Ting, J., Sunkin, S.M., Feng, D., Anastassiou, C.A., Barkan, E., et al. (2019). Classification of electrophysiological and morphological neuron types in the mouse visual cortex. Nat Neurosci 22, 1182–1195. 10.1038/s41593-019-0417-0.

69. Arkhipov, A., Gouwens, N.W., Billeh, Y.N., Gratiy, S., Iyer, R., Wei, Z., Xu, Z., Abbasi-Asl, R., Berg, J., Buice, M., et al. (2018). Visual physiology of the layer 4 cortical circuit in silico. PLoS Comput Biol 14, e1006535. 10.1371/journal.pcbi.1006535.

70. 70. Lorente De Nó, R. (1947). Action potential of the motoneurons of the hypoglossus nucleus. J. Cell. Comp. Physiol. 29, 207–287. 10.1002/jcp.1030290303.

71. 71. Teleńczuk, B., Dehghani, N., Le Van Quyen, M., Cash, S.S., Halgren, E., Hatsopoulos, N.G., and Destexhe, A. (2017). Local field potentials primarily reflect inhibitory neuron activity in human and monkey cortex. Sci Rep 7, 40211. 10.1038/srep40211.

72. Logothetis, N.K. (2002). The neural basis of the blood–oxygen–level–dependent functional magnetic resonance imaging signal. Phil. Trans. R. Soc. Lond. B 357, 1003–1037. 10.1098/rstb.2002.1114.

73. Buzsáki, G., and Draguhn, A. (2004). Neuronal Oscillations in Cortical Networks. Science 304, 1926– 1929. 10.1126/science.1099745.

74. Saleem, A.B., Lien, A.D., Krumin, M., Haider, B., Rosón, M.R., Ayaz, A., Reinhold, K., Busse, L., Carandini, M., and Harris, K.D. (2017). Subcortical Source and Modulation of the Narrowband Gamma Oscillation in Mouse Visual Cortex. Neuron 93, 315–322. 10.1016/j.neuron.2016.12.028.

75. Storchi, R., Bedford, R.A., Martial, F.P., Allen, A.E., Wynne, J., Montemurro, M.A., Petersen, R.S., and Lucas, R.J. (2017). Modulation of Fast Narrowband Oscillations in the Mouse Retina and dLGN According to Background Light Intensity. Neuron 93, 299–307. 10.1016/j.neuron.2016.12.027.

76. Mazzoni, A., Logothetis, N.K., and Panzeri, S. (2012). The information content of Local Field Potentials: experiments and models. Preprint at arXiv, 10.48550/ARXIV.1206.0560 10.48550/ARXIV.1206.0560.

77. Haider, B., Schulz, D.P.A., Häusser, M., and Carandini, M. (2016). Millisecond Coupling of Local Field Potentials to Synaptic Currents in the Awake Visual Cortex. Neuron 90, 35–42. 10.1016/j.neuron.2016.02.034.

78. Hwang, E.J., and Andersen, R.A. (2013). The utility of multichannel local field potentials for brain– machine interfaces. J. Neural Eng. 10, 046005. 10.1088/1741-2560/10/4/046005.

79. Bazelot, M., Dinocourt, C., Cohen, I., and Miles, R. (2010). Unitary inhibitory field potentials in the CA3 region of rat hippocampus. The Journal of Physiology 588, 2077–2090. 10.1113/jphysiol.2009.185918.

80. Haider, B., Häusser, M., and Carandini, M. (2013). Inhibition dominates sensory responses in the awake cortex. Nature 493, 97–100. 10.1038/nature11665.

81. Poulet, J.F.A., and Petersen, C.C.H. (2008). Internal brain state regulates membrane potential synchrony in barrel cortex of behaving mice. Nature 454, 881–885. 10.1038/nature07150.

82. Okun, M., Naim, A., and Lampl, I. (2010). The Subthreshold Relation between Cortical Local Field Potential and Neuronal Firing Unveiled by Intracellular Recordings in Awake Rats. J. Neurosci. 30, 4440– 4448. 10.1523/JNEUROSCI.5062-09.2010.

83. Anderson, J., Lampl, I., Reichova, I., Carandini, M., and Ferster, D. (2000). Stimulus dependence of two- state fluctuations of membrane potential in cat visual cortex. Nat Neurosci 3, 617–621. 10.1038/75797.

84. Cavallari, S., Panzeri, S., and Mazzoni, A. (2014). Comparison of the dynamics of neural interactions between current-based and conductance-based integrate-and-fire recurrent networks. Front. Neural Circuits 8. 10.3389/fncir.2014.00012.

85. Anderson, J.S., Carandini, M., and Ferster, D. (2000). Orientation Tuning of Input Conductance, Excitation, and Inhibition in Cat Primary Visual Cortex. Journal of Neurophysiology 84, 909–926. 10.1152/jn.2000.84.2.909.

86. Carandini, M., and Ferster, D. (2000). Membrane Potential and Firing Rate in Cat Primary Visual Cortex. J. Neurosci. 20, 470–484. 10.1523/JNEUROSCI.20-01-00470.2000.

87. Monier, C., Chavane, F., Baudot, P., Graham, L.J., and Frégnac, Y. (2003). Orientation and Direction Selectivity of Synaptic Inputs in Visual Cortical Neurons. Neuron 37, 663–680. 10.1016/S0896-6273(03)00064-3.

88. Priebe, N.J., and Ferster, D. (2005). Direction Selectivity of Excitation and Inhibition in Simple Cells of the Cat Primary Visual Cortex. Neuron 45, 133–145. 10.1016/j.neuron.2004.12.024.

89. Polack, P.-O., Friedman, J., and Golshani, P. (2013). Cellular mechanisms of brain state–dependent gain modulation in visual cortex. Nat Neurosci 16, 1331–1339. 10.1038/nn.3464.

90. Holcman, D., and Tsodyks, M. (2006). The Emergence of Up and Down States in Cortical Networks. PLoS Comput Biol 2, e23. 10.1371/journal.pcbi.0020023.

91. Wilson, C. (2008). Up and down states. Scholarpedia 3, 1410. 10.4249/scholarpedia.1410.

92. Campagnola, L., Seeman, S.C., Chartrand, T., Kim, L., Hoggarth, A., Gamlin, C., Ito, S., Trinh, J., Davoudian, P., Radaelli, C., et al. (2022). Local connectivity and synaptic dynamics in mouse and human neocortex. Science 375, eabj5861. 10.1126/science.abj5861.

93. Buchholz, M.O., Gastone Guilabert, A., Ehret, B., and Schuhknecht, G.F.P. (2023). How synaptic strength, short-term plasticity, and input synchrony contribute to neuronal spike output. PLoS Comput Biol 19, e1011046. 10.1371/journal.pcbi.1011046.

94. Moreni, G., Pennartz, C.M.A., and Mejias, J.F. (2023). Synaptic plasticity is required for oscillations in a V1 cortical column model with multiple interneuron types. Preprint, 10.1101/2023.08.27.555009 10.1101/2023.08.27.555009.

95. Spiess, R., George, R., Cook, M., and Diehl, P.U. (2016). Structural Plasticity Denoises Responses and Improves Learning Speed. Front. Comput. Neurosci. 10. 10.3389/fncom.2016.00093.

96. Gallinaro, J.V., and Rotter, S. (2018). Associative properties of structural plasticity based on firing rate homeostasis in recurrent neuronal networks. Sci Rep 8, 3754. 10.1038/s41598-018-22077-3.

97. Einevoll, G.T., Pettersen, K.H., Devor, A., Ulbert, I., Halgren, E., and Dale, A.M. (2007). Laminar Population Analysis: Estimating Firing Rates and Evoked Synaptic Activity From Multielectrode Recordings in Rat Barrel Cortex. Journal of Neurophysiology 97, 2174–2190. 10.1152/jn.00845.2006.

98. Dai, K., Gratiy, S.L., Billeh, Y.N., Xu, R., Cai, B., Cain, N., Rimehaug, A.E., Stasik, A.J., Einevoll, G.T., Mihalas, S., et al. (2020). Brain Modeling ToolKit: An open source software suite for multiscale modeling of brain circuits. PLoS Comput Biol 16, e1008386. 10.1371/journal.pcbi.1008386.

99. Wright, S. (1921). Correlation and causation. Journal of agricultural research 20.*7*: 557.

## Supplemental text references

1. Mazzoni, A., Lindén, H., Cuntz, H., Lansner, A., Panzeri, S., and Einevoll, G.T. (2015). Computing the Local Field Potential (LFP) from Integrate-and-Fire Network Models. PLoS Comput Biol 11, e1004584. 10.1371/journal.pcbi.1004584.

2. Teeter, C., Iyer, R., Menon, V., Gouwens, N., Feng, D., Berg, J., Szafer, A., Cain, N., Zeng, H., Hawrylycz, M., et al. (2018). Generalized leaky integrate-and-fire models classify multiple neuron types. Nat Commun 9, 709. 10.1038/s41467-017-02717-4.

